# A damaged-informed lung ventilator model for ventilator waveforms

**DOI:** 10.1101/2020.10.23.351320

**Authors:** Deepak K. Agrawal, Bradford J. Smith, Peter D. Sottile, David J. Albers

## Abstract

Motivated by desire to understand pulmonary physiology and pathophysiology, scientists have developed models of pulmonary physiology. However, pathophysiology and interactions between human lungs and ventilators, e.g., ventilator-induced lung injury (VILI), present problems for modeling efforts. Real-world injury is too complex for simple models to capture, and while complex models tend not to be estimable with clinical data, limiting both the clinical utility with existing approaches. To address this gap, we present a damaged-informed lung ventilator (DILV) model to model and quantify patient-ventilator interactions and lung health. This approach relies on systematically mathematizing the pathophysiologic knowledge clinicians use to interpret lung condition from ventilator waveform data. This is achieved by defining clinically relevant features in the ventilator waveform data that contain hypothesis-driven information about pulmonary physiology, patient-ventilator interaction, and ventilator settings. To capture these features, we develop a modelling framework where the model has enough flexibility to reproduce commonly observed variability in waveform data. We infer the model parameters with clinical (human) and laboratory (mouse) data. The DILV model can reproduce essential dynamics of differently damaged lungs for tightly controlled measurements in mice and uncontrolled human intensive care unit data in the absence and presence of respiratory effort. Estimated parameters correlate with known measures of lung physiology, including lung compliance. This method has the potential to translate laboratory physiology experiments to clinical applications, including pathways for high fidelity estimates of lung state and sources of VILI with an end goal of reducing the impact of VILI and acute respiratory distress syndrome.

## INTRODUCTION

Mechanical ventilation is a life-saving therapy for patients who are unable to perform gas exchange by breathing on their own. When used incorrectly, mechanical ventilation has the potential to worsen lung injury through barotrauma, volutrauma, and atelectrauma that are collectively referred to as ventilator-induced lung injury (VILI). Furthermore, when there is a mismatch between the patient’s respiratory demands and the mechanical breath, termed ventilator dyssynchrony (VD), VILI may be more severe and may lead to poor outcomes.^1–7^ Additionally, there are conditions or syndrome such as acute respiratory distress syndrome (ARDS) that carry a high mortality rate and may be exacerbated, or even caused, by VILI.^8–11^ Therefore, identifying lung-protective ventilation to reduce VILI both important and challenging because the oxygenation needs are often in opposition to safe ventilation, leading to a complex interplay between the underlying pulmonary pathophysiology, ventilator mechanics, and patient-ventilator interactions (respiratory effort).^12–15^ The current standard of care dictates a formulaic application of low tidal volumes to reduce overdistension and positive end-expiratory pressure to maintain patency. This approach reduces VILI but does not prevent it in all cases and is not personalized.^16–18^. While such protocols have provided measurable improvements in outcomes, the formulaic approach could potentially be improved through personalization of individual respiratory mechanics and respiratory effort.^19^

Modern mechanical ventilators produce data in the form of time-dependent pressure, volume, and flow waveforms that contain a wealth of information about respiratory mechanics, patient-ventilator interactions, and ventilator settings. These data can be used to troubleshoot and optimize mechanical ventilation.^20,21^ However, ventilator waveforms are typically analyzed heuristically by visual inspection and, therefore, the outcome of such an analysis is limited by individual expertise and experience. ^20,21^ A quantitative interpretation of these complex signals could increase diagnostic accuracy and repeatability while facilitating the application of personalized lung-protective ventilation. One simple example of waveform quantification that is currently used in clinical care is the driving pressure, which serves as a readout of both patient condition and ventilator settings.^22^ In the current study, we seek to develop a methodology to provide a more comprehensive quantitative description of lung injury severity from ventilator waveform data.

The analysis we present herein is a departure from traditional modeling methods that link measured pressures and flows through physiologically-based parameters, such as the well-recognized single-compartment model that lumps the spatially heterogeneous lung mechanical properties into single values of resistance and compliance.^23–27^ In the traditional models, the entirety of the pressure and volume dynamics emerge from the hypothesized physiological mechanics. Due to this straightforward formulation, the single-compartment model is computationally efficient but often not be able to reproduce all of the features in waveform data, such as patient respiratory effort because the model lacks the complexity to allow such complex dynamics to emerge. On the other hand, more complicated formulations, including multi-compartment models, use many states and parameters that cannot be directly measured, such as recruitment pressure distributions, causing identifiability problems where there is no unique solution, or more often no convergent solution for parameter values. As such, those models require expensive data to estimate, data not available in humans, and require substantial computational resources. Even then, complex multi-compartment models often cannot produce all the relevant features present in the pressure and volume waveform data.^28–36^ One of the objectives is to identify, quantify the severity of, and understand the damage due to VILI and VD. This is out of reach with current fully mechanistic models, and while machine learning approaches can potentially be used to detect VD,^37^ they are not hypothesis-driven and therefore while these methods produce machinery that is explainable in the sense of identifying predictive sources of the classification task of VD/no-VD breaths, these methods are not compatible with physiology-anchored explanations of, e.g. VD.

Our approach offers the potential to overcome these limitations and provides both identifiability and fidelity by using mathematical models with interpretable parameters to recapitulate pressure and volume signals. This high fidelity is due, in part, to the limited dependence between the pressure and volume models. The relationship between components of the pressure and volume waveforms are used to define specific physiologic features, just as the quasi-static compliance is defined from the observed ratio of tidal volume and driving pressure. We anticipate that this approach will find applications in real-time clinical readouts of ventilation safety, long-term monitoring to detect changes in patient condition, and as a quantitative outcome measure for clinical trials.

Here we build a model that takes as baseline “healthy” breaths, and then adds terms that correspond to deviations from healthy breaths whose hypothesized source include VILI/VD/pathophysiological features observed in ventilator waveforms. By estimating the model, we identify the presence and severity of deviations from normal in a way that has a physiologically-based hypothesis attached to it. In essence, we compute hypothesis-driven breath phenotypes that may potentially be used to run trials on outcomes related to the frequency and strength of certain deviations that correspond to types of VD. One primary goal of this work is to tie these phenotypes to the pathogenesis of VILI and the associated cause of VILI. To this end, we work to understand what damage is happening to the lungs, and under what circumstances, by estimating the model for both humans and mice where (1) the waveforms are very similar and only differ in scale and more importantly, (2) under the same hypothesized conditions - created in the mice and observed in the humans and believed to be present. In this case, we can dig deeply into the mouse lungs in a way not possible with humans and carefully and deeply characterize the nature, severity of lung damage. In these ways, this modeling framework is meant to be a bridge between the clinic and laboratory.

In the current study, we first identify clinically important deviant features in typical volume and pressure waveform data. We then separately define the models for volume and pressure waveforms as the sum of a set of terms through which we modularly capture physiologically relevant features. This approach allows independent modeling of the components of damage so that clinical and physiologic knowledge can be used to constrain the model. We named this model the damage-informed lung-ventilator (DILV) as it contains information about both lung physiology and ventilator dynamics. To demonstrate the model’s flexibility, the volume and pressure models are qualitatively validated in a simulation study where we show various relevant features that are commonly observed in health and disease. We then identify the parameters that correspond to interpretable pathophysiology by analyzing simulated pressure-volume waveforms (also known as PV loops) and by relating the changes in the DILV model with known lung compliance and resistance. Finally, in a quantitative verification, we demonstrate that the model can accurately and uniquely represent laboratory and clinical ventilator data, which includes mouse model and human-intensive care unit (ICU) ventilator data in the absence and presence of respiratory effort.^37^ Through a comprehensive comparison between the DILV model and the single-compartment model, we demonstrate that our approach can produce the features present in the waveform data and report values of lung compliance. Temporal changes in the model parameters are compared to other assessments of injury severity and qualitative features of the pressure and volume waveforms.

## METHODS

### Identifying relevant and realistic variables for the model

Our goal is to develop a model that can reproduce most common physiologically-relevant features present in waveforms data so that the model can identify, understand, and quantify lung pathophysiology, particularly VILI, in clinical settings. Therefore, it is critical to identify the appropriate complexity of the model and anchor the model to hypothesized lung/ventilator/damage mechanics to achieve the desired outcome. This paper represents a starting point for this approach, but the work is far from complete. The model presented here does not represent all types of VILI, VD, and pathophysiology. Instead, as a methodological starting point, the DILV model can represent some types of VILI, VD, and pathophysiology. The model construction shows a pathway by adding model features tied to particular kinds of VILI or hypothesized pathophysiology, allowing the model to include a wide swath of VILI and VD eventually.

To start, mechanical ventilation is characterized by three measured state variables which vary over time: volume, pressure and flow. In human ventilator data these time-dependent waveforms have diverse features arising from physiology, pathophysiology, the ventilator, and health care process effects such as clinical interventions, and patient-ventilator interaction. ^39–41^ The flow is the time-derivative of volume and so the volume variable contains the same information about the underlying lung mechanics but in a different representation.^34^ In this study, we focused on two state-variables, the volume and pressure.

In the simplest ventilation mode, there is one variable that is primarily controlled by the ventilator, e.g., pressure or volume, while the other variable, e.g., volume or pressure, is free to vary, referred to as pressure-controlled ventilation (PCV) or volume-controlled (VCV) respectively. In this case, only the ‘free’ variable contains direct information about the respiratory mechanics of the patient.^34,43^ Moreover, in some models there is a rigid coupling between the controlled and free states that often limits the model flexibility, precluding the model from reproducing some features that are present in the clinical data. For example, the single-compartment model performs a linear transformation between pressure and volume variables due to the fixed coupling defined as the sum of linear resistive and elastic contributions. ^25,26,34^ We, therefore, do not explicitly couple the controlled (also known as an independent) and free (also known as a dependent) variables such that the volume and pressure models will be independent of one another. Human ventilators also have an expansive set of other modes, the most notable of which are the patient-triggered modes where the ventilator’s action is triggered by patients such as inspiratory effort. These modes can be very lung protective, but they can also lead to complex forms of VD that are difficult to model. Patient-triggered modes are the most commonly used modes for humans unless the human is given neuromuscular blockades.^42^

### Identifying important features in the volume and pressure waveform

In PCV, the pressure is the controlled variable and is set by the ventilator, shown in blue in Fig. 1a. ^20,21^ In this case, the volume is a free variable and generally has two characteristic features that can be divided into two features, shown in red in Fig. 1a. The first feature is the inspiration, denoted as A in Fig. 1a, which continues until the amount of gas delivered in that breath is reached (the tidal volume). The pressure and lung elastic recoil are at equilibrium. The second feature is expiration, denoted as B in Fig. 1a. Depending on the ventilator settings and lung condition, the gradient of the rising and falling signals can vary across patients and in the same patient over time. Therefore, the model must be able to represent these features independently. Accordingly, the gradients of inspiration and expiration of volume are features that are variable and estimable in the volume model.

**Figure 1:**
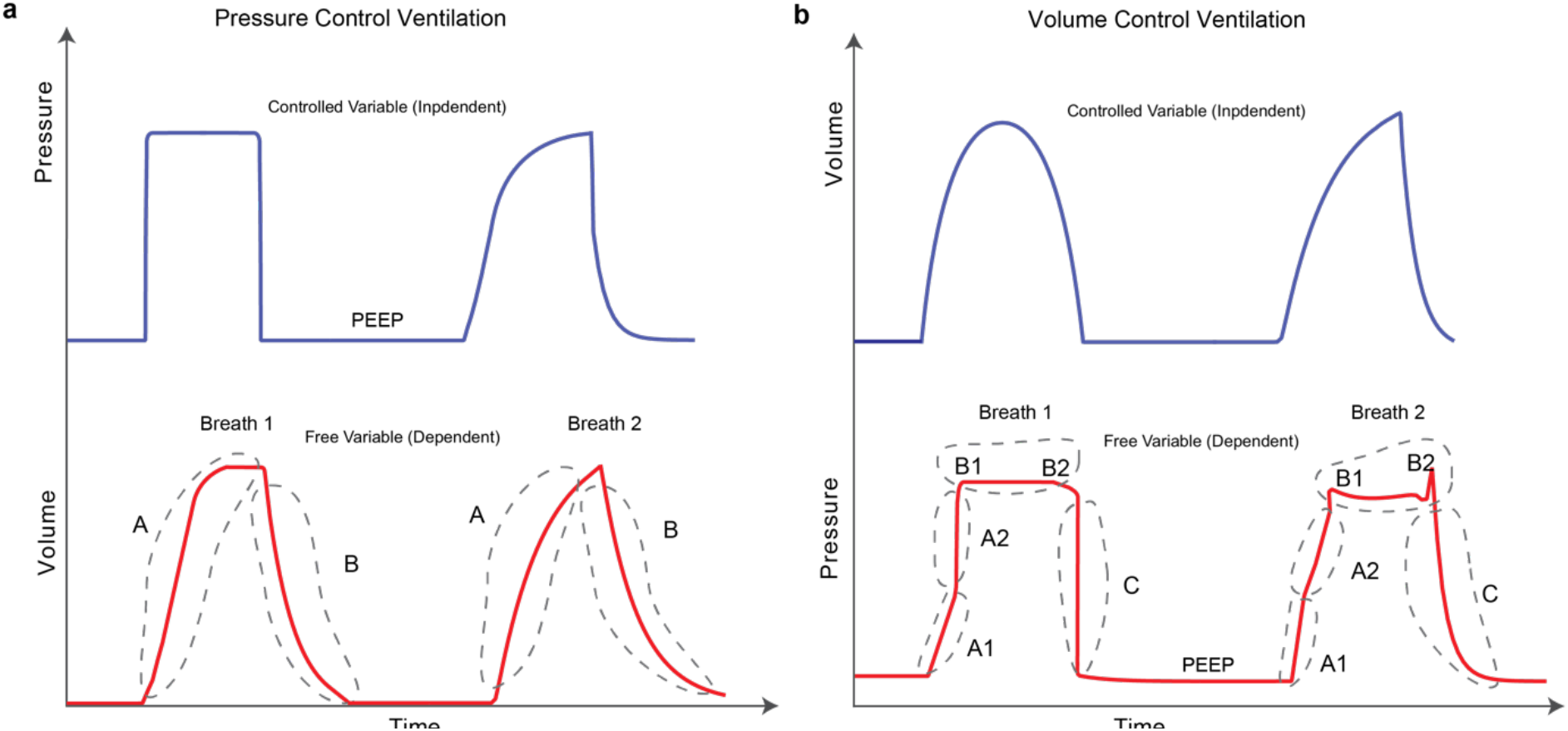
Graphical representation of typical volume and pressure waveforms. **(a)** In PCV, the volume is a free (also known as a dependent) variable and generally has two distinct features. The rising and falling of the volume signal during inspiration and expiration, respectively, are denoted as A and B. **(b)** When pressure is a free variable (in VCV), there can be multiple features in the waveform that contain useful information. The gradient of the rising signal in which pressure continues to increase during inspiration can have two distinct features, denoted as A1 and A2. These two features define the gradient of the rising signal before and after the inflection point such that there may be abrupt increases (breath 1) or decreases (breath 2) in the signal gradient. The shape of the plateau pressure is captured using features B1 and B2 such that there may be a peak at the beginning (B1-breath 2) and/or at the end (B2-breath 2) of the plateau. Finally, the gradient of the falling signal is captured using feature C that represents the expiration process. The baseline pressure is known as positive end-expiratory pressure (PEEP), and often used in ARDS patients to maintain an open lung.^45^ Here, the controlled and free variables are shown in the blue and red colors, respectively.

The characteristic shape of the pressure waveform can vary more dramatically than the volume waveform (Fig. 1b). When pressure is a free variable, such as in VCV, the pressure waveform has several important features that convey information about lung condition and ventilator-patient interaction, shown in red in Fig. 1b. Based on clinical understanding, we identified five important features in the pressure waveform that may be affected by lung condition. Features A1 and A2 in Fig. 1b determine the gradient of the inspiration. The time-varying graph of inspiration can have two distinct modes where the gradient of the signal may increase (Fig. 1b, breath 1) or decrease (Fig. 1b, breath 2) during inspiration. These features are hypothesized to correspond to the volume-dependent decrease in lung compliance (breath 1) or an increase in compliance due to recruitment (breath 2). ^25,26^ Features B1 and B2 (Fig. 1b) are related to the shape of the waveform at the start and end of the plateau pressure, which is a period of constant pressure. There may be peaks at the beginning (B1) and/or at the end (B2) of the plateau pressure, which are hypothesized to correspond to inspiratory flow resistance and patient effort, respectively. ^20,34^ Feature C in Fig. 1b corresponds to the gradient of expiration. We also model the constant baseline pressure, known as the positive end-expiratory pressure (PEEP), because it is a key independent variable in ARDS management.^44,45^ Note that in hybrid ventilation modes, there may be scenarios where both pressure and volume variables are partially controlled and so, in those cases, both the waveforms can be confounded in additionally complex ways and would admit more subtle interpretations.

### Constructing the damage-informed lung ventilator model

#### Construction of the volume model

Irrespective of the state variable, the models have periodic dynamics with a frequency defined by the respiratory rate (breaths/min) that should be the same in pressure and volume waveform models. In addition to this constraint, the volume model has two additional features, the rate of inspiration and expiration (A and B in Fig. 1a, respectively). Volume model development begins by modeling the respiratory rate with a sinusoidal function (*f_s1_*):

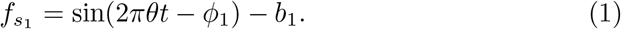

Here, the respiratory frequency (breaths/s) is set by *θ* and *t* represents time in seconds while parameter *ϕ_1_* allows to control the starting point in the respiratory cycle. To control the rate of inspiration or expiration while maintaining the periodicity, we create a periodic rectangular waveform function *f_b1_* by combining the sinusoidal function with hyperbolic tangent function:

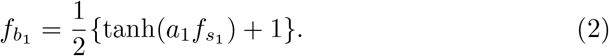

To control the smoothness of the rectangular waveform, we added a smoothing parameter *a_1_*. The other terms (1/2, +1) are added to generate a rectangular waveform that has a zero-base value and unit amplitude. To control the duty cycle of the rectangular waveform that sets inspiratory:expiratory (**I:E**) ratio, we used parameter *b_1_* such that the zero value of *b_1_* corresponds to 1:1 I:E ratio.

Fig. 1a shows additional model features: the rate of inspiration (A) and expiration (B). To represent these rates independently, we define the volume (*V*) using the rectangular waveform as a base waveform:

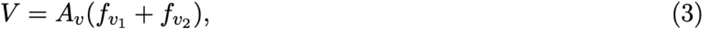

Where *f_v1_* term produces the inspiration part of the volume signal (feature A):

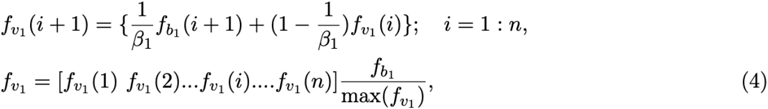

and *f_v2_* term produces the expiration part of the volume signal (feature B):

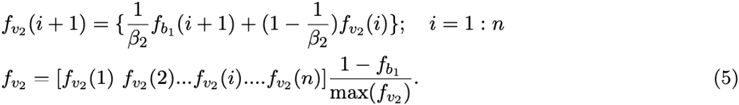

Here, *β_1_* and *β_2_* control the gradient of the inspiration and expiration, respectively, while *A_v_* controls the amplitude of the volume waveform. Figure 2a shows the volume waveform (top plot) and the constitutive terms added through with Eqns. 1-5.

**Figure 2:**
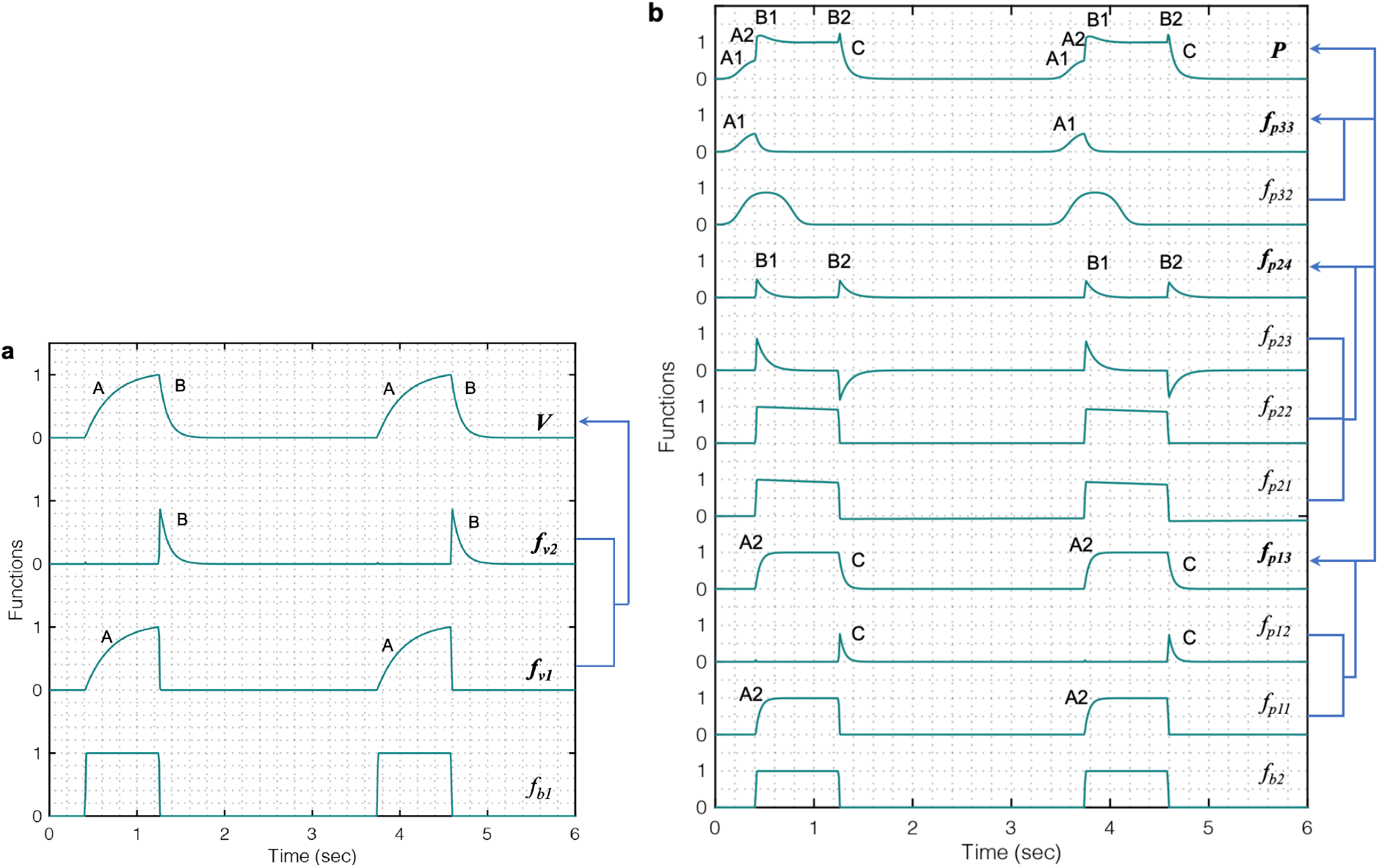
Simulated response of various terms that make up the damage-informed volume (*V*) and pressure model (*P*). **(a)** A periodic rectangular waveform *f_b1_* is used to create terms *f_v1_* and *f_v2_* through which the gradient of the rising (feature A) and falling (feature B) signals in the volume waveform are controlled, respectively. Equations 1-5 were used to simulate the response within each term with parameter values *θ* = 0.3, *a_1_* = 200, *b_1_* = 0.7, *ϕ_1_* = 0, *β_1_* = 30, *β_2_* = 10, *A_v_* = 1. **(b)** A periodic rectangular waveform (*f_b2_*) serves as a basis of the pressure model. The overall shape of the pressure waveform, which defines the gradient of the inspiration and expiration signals, is formed using *f_p13_* comprised of the rising signal of *f_p11_* (A2) and falling signal of *f_p12_* (C). The shape of the plateau pressure is defined by *f_p24_*, where the output of *f_b2_* is processed via *f_p21_*, *f_p22_* and *f_p23_* to produce peaks at the beginning (B1) and end (B2) of the plateau pressure. The shape of the rising signal at low volume (A1) is defined by *f_p33_*, where a short pulse is produced via *f_p31_* and reshaped via *f_p32_*. Note that the amplitude terms *A_p1_*, *A_p2_* and *A_p3_* control the amplitude of *f_p13_*, *f_p24_* and *f_p33_*, respectively. Equations 6-18 were used to simulate the response of each term with parameter values *θ* = 0.3, *a_2_* = 200, *b_2_* = 0.7, *ϕ*_2_ = 0, *a_3_* = 10, *b_3_* = 0.9, *ϕ*_3_ = −0.6, *β_3_* = *β_4_* = 5, *β_5_* = 1.001, *β_6_* = 1.1111, *A_p1_* = 1, *A_p2_* = 0.5, *A_p3_* = 0.5, *A_p4_* = 0. Note that the model variability shown here is independent of the ventilator mode.

#### Construction of the pressure model

The pressure model has five explicit features that might correspond to lung pathophysiology and ventilator settings. These features are depicted in Fig. 1b. Features A1 and A2 capture the gradient of the rising signal during inspiration at low (A1) and high (A2) volume and might be correlated with lung compliance. Features B1 and B2 capture the shape of the peaks at the beginning (B1) and end (B2) of the plateau pressure and might reflect changes in inspiratory flow resistance and patient effort, respectively. Finally, feature C captures the rate of change of the pressure during expiration. To build a model that can capture all these features and be able to estimate parameters effectively, we used a specific set of parameters that were used to build the components that contribute to the pressure model modularly.

The pressure model construction begins like the volume model, with a sinusoid. Because volume and pressure are coupled through their period, we enforce this constraint by requiring that both models have the same respiratory frequency (*θ*) in their base periodic sinusoid:

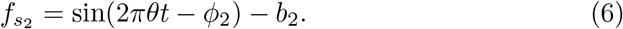

Because the pressure may lag or lead the volume, we include a phase shift term, *ϕ*_2_ in the sinusoid. To account for additional variations in the duty cycle of the rectangular waveform, we added the parameter *b_2_* that defines the **I:E** ratio. We then create a rectangular waveform *f_b2_* as we did for the volume model using the hyperbolic tangent, or:

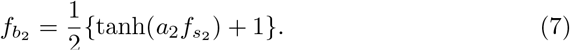

The smoothness of the rectangular waveform is controlled via the parameter *a_2_*. The five key features in pressure are represented with three additional terms: (i) *f_p13_* defines the rates of pressure change during inspiration and expiration, (ii) *f_p24_* determines the peaks at the beginning and end of the pressure plateau, and (iii) *f_p33_* specifies the gradient of the initial rising signal during inspiration, leaving us with the full the pressure model (P):

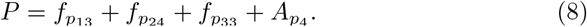

The constant parameter *A_p4_* corresponds to the baseline pressure value (PEEP). The rates of pressure change during inspiration and expiration (see A2, and C in Fig. 1b, respectively) are:

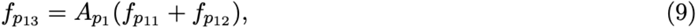

where *f_p11_* term produces the rising part of the pressure signal:

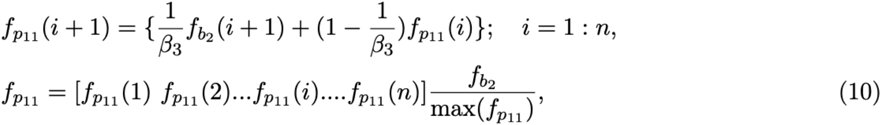

and *f_p12_* term produces the falling part of the pressure signal:

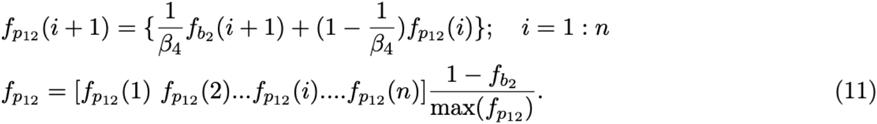

Here, *β_3_* and *β_4_* control the gradient during inspiration and expiration, respectively. The next set of features, the peaks at the beginning and end of plateau pressure (B1 and B2 in Fig. 1b), are represented by:

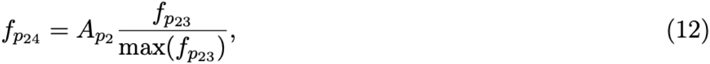

where *f_p21_* and *f_p22_* terms create the initial shape of peaks at the plateau pressure:

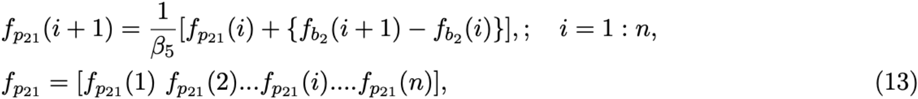

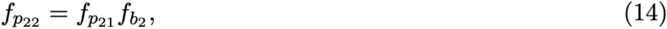

and *f_p32_* term further reshapes both the peaks:

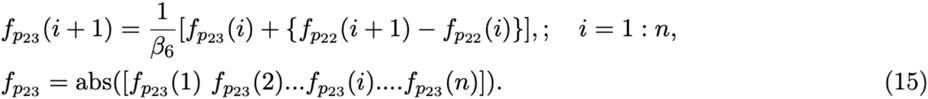

The parameters *β_5_* and *β_6_* control the shape of both the peaks, which are present at the plateau pressure. Finally, the gradient of the initial rate of inspiration (Fig. 1b, A1) is modeled by:

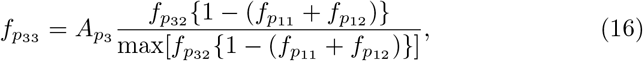

where a short pulse is produced via *f_p31_*:

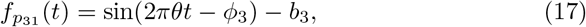

and reshaped via *f_p32_* term:

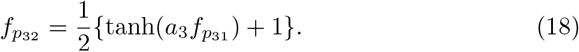

The position, shape and gradient of the rising signal, produced by *f_p33_* term are controlled using the parameters *ϕ_3_*, *b_3_* and *a_3_*, respectively. Figure 2b shows the pressure waveform and the constitutive terms added through Eqns. 6-18. The pressure waveform (top plot) composed of the three terms *f_p13_*, *f_p24_* and *f_p33_* that capture the gradient of the inspiration (A2) and expiration signals (C), the shape of the plateau pressure (B1 and B2), and the shape of the rising signal at low volume (A1), respectively.

### Qualitative model validation, parametric descriptions and simulated model variability

The first step in model validation—qualitative validation^38^—involves demonstrating the model has enough flexibility to recapitulate the key features that are often seen in clinically collected volume and pressure waveform data.

#### Volume model flexibility

The volume model flexibility is demonstrated in Fig. 3 where we vary the rates of inspiration (feature A) and expiration (feature B) through the terms *f_v1_* and *f_v2_*, which is controlled by parameters *β_1_* and *β_2_* (Fig. 3a), respectively. The full variability of terms *f_v1_* and *f_v2_* is shown in Supplementary Fig. S1 (all Supplemental Material is available at https://www.biorxiv.org/content/10.1101/2020.10.23.351320v1.supplementary-material). Additionally, the amplitude of the volume waveform is controlled by *A_v_* (Fig. 3c), variations in respiratory rate are controlled by *θ* (Fig. 3d), and the I:E ratio is altered through the parameter *b_1_* that changes the duty cycle of the rectangular base waveform (Fig. 3e). Finally, the starting point of the breath in the breathing cycle and the smoothness of the volume waveform are set by *ϕ_1_* and *a_1_* as shown in Fig. 3f and Supplementary Fig. S2a,b, respectively.

**Figure 3:**
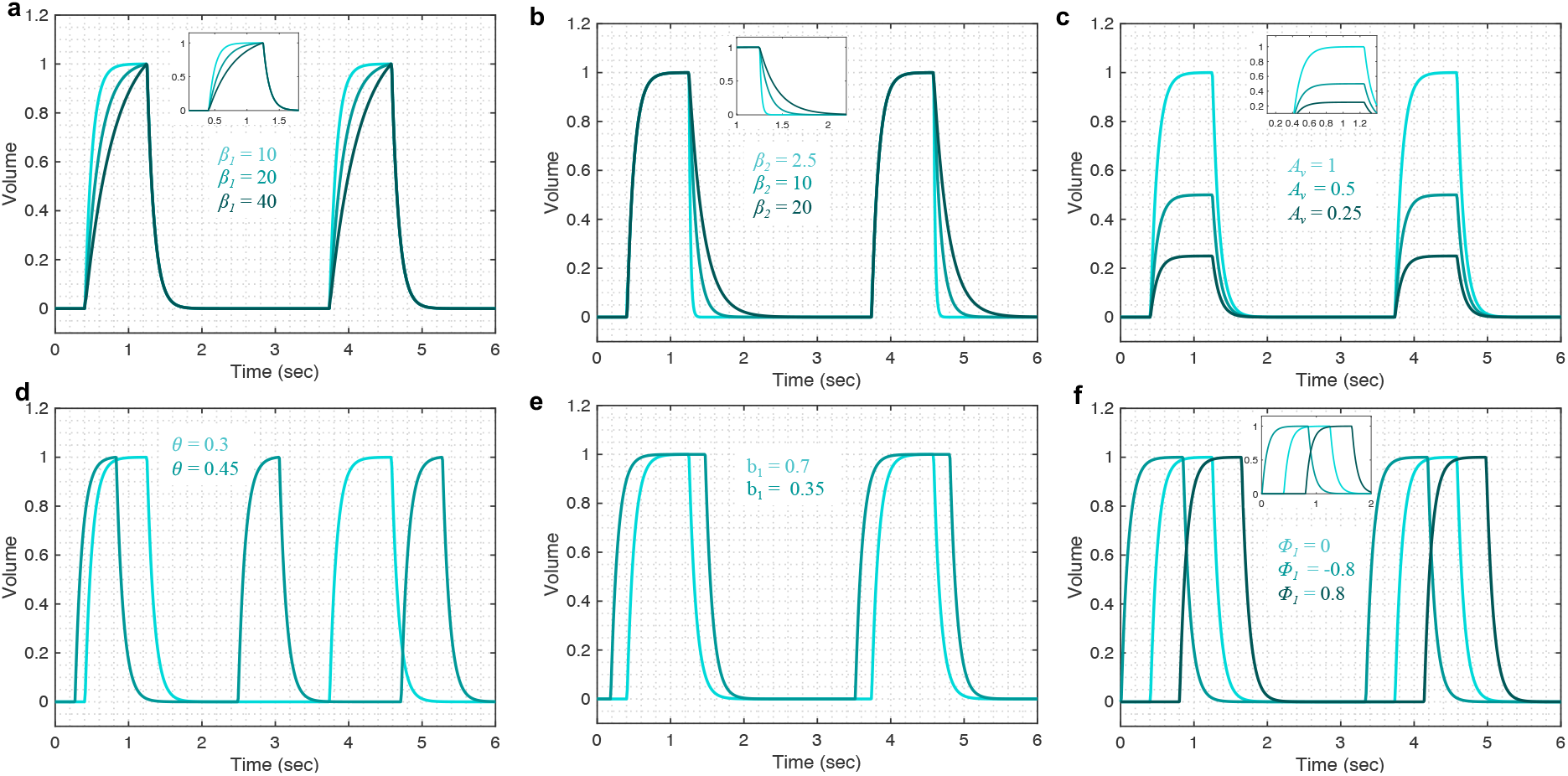
Demonstrating the volume model flexibility by varying parameters that alter characteristic features of the volume waveform. The gradient of the rising and falling signals can be altered using the **(a)** *β_1_* and **(b)***β_2_* parameters, respectively. Increased values of these parameters increase the transient time for the signal to reach the same volume level. **(c)** The amplitude of the waveform can be altered using the parameter *A_v_*. **(d)** Changes in the respiratory frequency (*θ*) change the period of the breath while **(e)** the I:E ratio (inspiratory to expiratory time ratio) can be modified using the *b_1_* parameter. **(f)** The starting point of the inspiration can be modified using the parameter *ϕ_1_*. The output of the model (*V*) was calculated using Eqns. 1-5 while considering *θ* = 0.3, *a_1_* = 200, *b_1_* = 0.7, *ϕ_1_* = 0, *β_1_* = 30, *β_2_* = 10, *A_v_* = 1. The respective variation in the functions that make the volume model is shown in Supplementary Fig. S1 for each case. Note that the model variability shown here is independent of the ventilator mode.

#### Pressure model flexibility

The pressure model flexibility is demonstrated in Fig. 4 where we vary the five features of the pressure waveform via respective parameters: variation in the rate of change of the pressure before (A1 in Fig. 2b) and after (A2 in Fig. 2b) the inflection point during inspiration; the shape of the peaks at the beginning (B1 in Fig. 2b) and end (B2 in Fig. 2b) of the plateau pressure; and variation in the rate of change of the pressure during expiration (C in Fig. 2b). In brief, these features are controlled by the following parameters.

**Figure 4:**
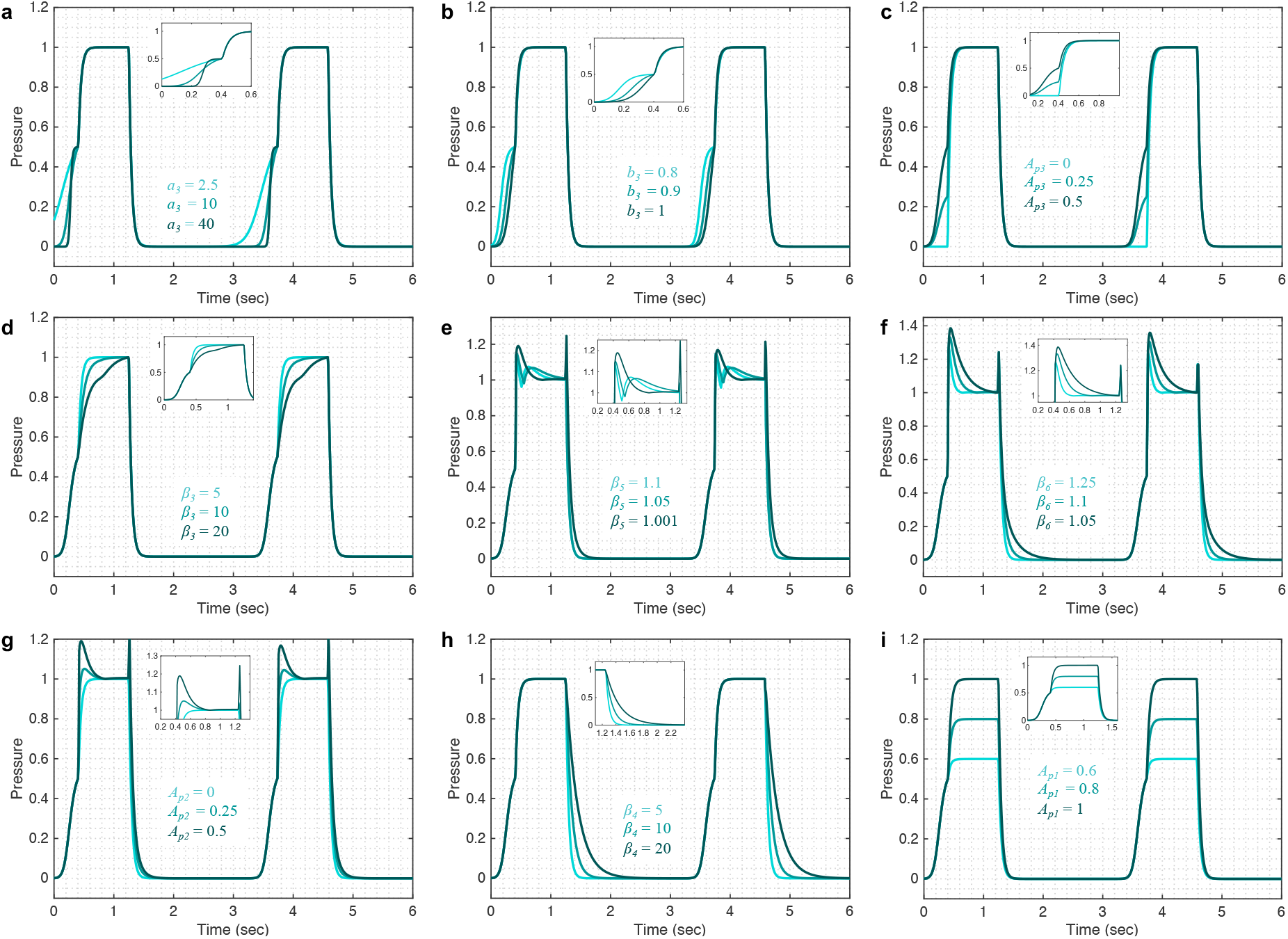
Demonstrating the pressure model flexibility by altering relevant features in the pressure waveform. The initial gradient of the pressure signal during inspiration at low volume (A1) is controlled by **(a)** the *a_3_* and **(b)***b_3_* parameters while **(c)** the inflection point location can be altered using the *A_p3_* parameter. **(d)** The gradient of the rising signal after the inflection point (A2), is controlled by the *β_3_* parameter. **(e)** The shapes of the peaks at the beginning (B1) and at the end (B2) of the plateau are regulated by the *β_5_* parameter when *A_p4_* = 0.5. **(f)** The sharpness of these peaks can be altered further by the *β_6_* parameter for a given shape while **(g)** the amplitude of the peaks is controlled by the *A_p2_* parameter. **(h)** The gradient of the falling signal (C) during expiration can be modified by the *β_4_* parameter. **(i)** The amplitude of the waveform can be altered using the parameter *A_p1_*. Equations 6-18 were used to simulate the response of the pressure model while considering *θ* = 0.3, *a_2_* = 200, *b_2_* = 0.7, *ϕ*_2_ = 0, *a_3_* = 10, *b_3_* = 0.9, *ϕ*_3_ = −0.6, *β_3_* = *β_4_* = 5, *β_5_* = 1.001, *β_6_* = 1.1111, *A_p1_* = 1, *A_p2_* = 0.5, *A_p3_* = 0.5, *A_p4_* = 0. The respective variations in the terms that make the pressure model are shown in the Supplementary Fig. S3 for each case. Note that the model variability shown here is independent of the ventilator mode.

The initial gradient of the pressure during inspiration (A1) is controlled by the *a_3_* parameter such that higher values of *a_3_* result in a slower rising signal (Fig. 4a). The full variation these terms are capable of is shown in Supplementary Fig. S3. The shape of the initial gradient signal (A1) before inflection point can be altered using the *b_3_* parameter (Fig. 4b) and the amplitude of the initial gradient alteration is controlled by the *A_p3_* parameter (Fig. 4c). By setting *A_p3_* parameter to zero, feature A1 can be removed from the pressure waveforms. The rate of inspiratory pressure after the inflection point (A2) is specified by *β_3_* such that higher values of *β_3_* result in a slower rising signal (Fig. 4d). The shapes of the peaks at the beginning (B1) and end (B2) of the plateau pressure are controlled by several parameters. The overall shape of the peaks is controlled by the *β_5_* (Fig. 4e) and the sharpness of these peaks can be further altered by *β_6_* (Fig. 4f). The amplitude of the peaks is controlled by *A_p2_* (Fig. 4g). By setting *A_p2_* parameter to zero, features B1 and B2 can be turned off. Additional control of features B1 and B2 can be achieved in combination with parameter *β_3_* shown in Supplementary Fig. S2c-f. The rate of pressure decrease during expiration (C) is specified by *β_4_* such that higher values of *β_4_* result in a slower falling signal (Fig. 4h). Finally, the amplitude of the plateau pressure can be altered using the *A_p1_* parameter (Fig. 4i). The I:E ratio is defined by the *b_2_* parameter in the same way that parameter *b_1_* controls the I:E ratio in the volume model (Fig. 3e). A summary of model parameters is provided in Table 1.

**Table 1:** Interpreting damaged-informed lung ventilator model parameters. The parameters that are correlated with known measures of lung physiology are in bold.

| Parameter | Model relevance | Physiological relevance<br>(with increased values) |
| --- | --- | --- |
| **Volume model** |  |  |
| $\theta$ | Number of breaths/s. Higher values result in a higher number of breaths/s. | - |
| $a_1$ | Smoothness of the square waveform ($f_{b1}$). Higher values result in a sharper transition. | - |
| $b_1$ | I:E ratio. Higher values result in smaller inspiration cycle. | - |
| $\Phi_1$ | Starting of the inspiration point. Higher value results in a more delayed response. | - |
| $\beta_1$ | Gradient of the rising signal. Higher values result in a slower rising signal. | Lower compliance and/or higher resistance. |
| $\beta_2$ | Gradient of the falling signal. Higher values result in a slower falling signal | Higher expiration time constant. |
| $A_v$ | Peak amplitude | Higher overall compliance. |
| **Pressure model** |  |  |
| $b_2$ | I:E ratio. Higher values result in a smaller inspiration cycle. | - |
| $a_2$ | Smoothness of the square waveform. Higher values result in a sharper transition. | - |
| $\Phi_2$ | Starting of the $f_{b2}$ function. Higher value results in a more delayed response | - |
| **$a_3, b_3$** | Gradient of the pressure signal at low volume. Higher values result in a slower rise. | Higher low volume compliance. |
| $\Phi_3$ | Starting of the inspiration point. Higher value results in a more delayed response | - |
| $\beta_3$ | Gradient of the rising signal after inflection point. Higher values result in a slower rising signal | Higher high-volume compliance. |
| $\beta_4$ | Gradient of the falling signal. Higher values result in a slower falling signal. | - |
| $\beta_5, \beta_6$ | Shape of plateau. Higher value means a sharper peak | Associated with ventilator desynchrony. |
| $A_{p1}$ | Amplitude at plateau | Lower high-volume compliance. |
| $A_{p2}$ | Amplitude of the peaks at the plateau | |
| $A_{p3}$ | Higher values increase the amplitude of $f_{p33}$ function | Moves upper inflection point up |
| $A_{p4}$ | Pressure base line value | PEEP |

### Explicit physiological lung function hypothesized to model parameters to lung function

The model developed here is anchored to physiology through variations or deviations from nominal breath waveforms that are hypothesized to directly relate to VILI, VD, and pathophysiology observed in mechanical ventilation data collected in clinical settings. Throughout the paper, we list the physiological interpretations of the parameters - a short description of how the model parameters contribute to the model is provided in Table 1 - but here we will go into interpretative depth regarding the qualitative correlation between model parameters and the fundamental characteristics of the lung such as compliance and resistance. In this qualitative interpretation we consider only one variable (volume in PCV and pressure in VCV) while assuming the other waveform does not change breath-to-breath (pressure in PCV and volume in VCV).

We first show how the changes in the volume model parameters can be qualitatively related to changes in lung compliance and resistance when the volume variable is free. For that, we focus on three model parameters that might have direct physiological meaning: *β_1_*, *β_2_*, and *A_v_*. First parameter, *β_1_* might be inversely correlated with lung compliance as higher values of *β_1_* result in a lower inspiratory flow rate (Fig. 3a, Supplementary Fig. S4a). During PCV, the inspiratory flow rate will decrease with reduced compliance or increased resistance. A second parameter, *β_2_*, controls the gradient of expiration and is captured as feature B in Fig. 1a. Higher values of *β_2_* result in a longer expiration (cf. Fig. 3b, Supplementary Fig. S4a) and so *β_2_* is directly proportional to the expiratory time constant, which is the product of resistance and compliance. Therefore, an increase in parameter *β_2_* might suggest an increase in lung compliance. Finally, parameter *A_v_* controls the amplitude of the volume waveform and for the same pressure waveform (in PCV) indicate a direct correlation with compliance (Fig. 3c, Supplementary Fig. S4a). In VCV, parameter *A_v_* would present the tidal volume, which is set by the ventilator.

The pressure model has five parameters that are associated with aspects of lung compliance during VCV: *a_3_*, b_*3*_, *β_3_*, *A_p1_* and *A_p3_*. The parameter *a_3_*, which controls feature A1 (Fig. 4a), may be directly correlated with low-volume compliance as higher values of *a_3_* result in slower pressure rise at low volume while maintaining the shape of the gradient (Supplementary Fig. S4b). Additionally, parameter *b_3_* can also be used to control feature A1 (Fig. 4b, Supplementary Fig. S4b) and is directly related to the low-volume compliance. A third parameter, *β_3_* controls the rate of pressure increase above the inspiratory inflection point (A2), and higher values of *β_3_* result in slower pressure increase (Fig. 4d, Supplementary Fig. S4b), indicating *β_3_* might be correlated with high-volume compliance during VCV. A fourth parameter, *A_p1_,* defines the plateau pressure with higher values of *A_p1_* yielding higher plateau pressures (Fig. 4i, Supplementary Fig. S4b), indicating an inverse correlation between *A_p1_* and compliance during VCV. Finally, change in the upper inflection point (UIP) can be directly related to the *A_p3_* parameter such that higher values of *A_p3_* increase the UIP pressure as shown in Supplementary Fig. S4b. During PCV, these (and other) parameters may be directly controlled via a ventilator.

It is important to note that these interpretations are qualitative and valid only when a change is observed in one of the variables (volume or pressure) while the other waveform (pressure or volume) is held fixed. In cases where both the volume and pressure waveforms change simultaneously, additional interpretation is needed to establish the relationships between pressure and volume parameters. For example, when there is a change in the amplitude of volume and pressure simultaneously, we use the *A_v_*/*A_p1_* to assess compliance.

### Mouse Mechanical Ventilation Experiments

Three 8 to 10-week-old female BALB/c mice (Jackson Laboratories, Bar Harbor, ME, USA) were studied under University of Colorado, Anschutz Medical Campus Institutional Animal Care and used Committee (IACUC)-approved protocol (#00230). Anesthesia was induced with an intraperitoneal (IP) injection of 100 mg/kg Ketamine and 16 mg/kg Xylazine, a tracheostomy was performed with a blunted 18 ga metal cannula, and ventilation was started on the flexiVent small animal ventilator (SCIREQ, Montreal, QC, Canada). Anesthesia was maintained with 50 mg/kg Ketamine or 50 mg/kg Ketamine with 8 mg/kg Xylazine at 30 min intervals along with 50 μL IP 5% dextrose lactated Ringer’s solution. Heart rate was monitored via electrocardiogram.

Baseline ventilation, consisting of a tidal volume (Vt) = 6 ml/kg, PEEP = 3 cmH_2_O, and respiratory rate (RR) = 250 breaths/min, was applied for a 10 min stabilization period with recruitment maneuvers at 3 min intervals. Pressure and volume were recorded with a custom flowmeter based on our previously published design.^46^ Four types of ventilation were recorded for analysis: 1) VCV-PEEP0, consisting the baseline ventilation with PEEP = 0 cmH_2_O, 2) VCV-PEEP12 that was the baseline ventilation with PEEP = 12 cmH_2_O, 3) HighPressure-PEEP0 that consisted of a inspiratory pressure (Pplat) = 35 cmH_2_O and PEEP = 0 cmH_2_O with RR = 60 breaths/min, and 4) PCV-PEEP0 with Pplat = 10 and PEEP = 0 cmH_2_O with RR=70 breaths/min. After the initial measurements of the healthy lung, lung injury was induced with a 0.15 ml lavage with warm saline.^23^ This fluid was pushed into the lung with an additional 0.3 ml air, and suction was applied to the tracheal cannula with an approximate return of 0.05 ml. The mouse was then ventilated for 10 mins with the HighPressure-PEEP0 settings. The sequence of four measurement ventilation patterns (above) was repeated, then the mouse received 0.8 mg/kg IP pancuronium bromide to suppress respiratory efforts, and the measurements were repeated again.

### Human Data Collection

Between June 2014 and January 2017, 140 adult patients admitted to the University of Colorado Hospital medical intensive care unit (MICU) at risk for or with ARDS and requiring mechanical ventilation were enrolled within 12 hours of intubation.^47^ At risk patients were defined as intubated patients with hypoxemia and a mechanism of lung injury known to cause ARDS, who had not yet met chest x-ray or oxygenation criteria for ARDS. To facilitate the capture of continuous ventilator data, only patients ventilated with a Hamilton G5 ventilator were included. Patients requiring mechanical ventilation only for asthma, COPD, heart failure, or airway protection were excluded. Additionally, patients less than 18 years of age, pregnant, or imprisoned were excluded. The University of Colorado Hospital utilizes a ventilator protocol that incorporated the ARDS network low tidal volume protocol with the low PEEP titration table. The Colorado Multiple Institutional Review Board approved this study and waived the need for informed consent.

Baseline patient information including age, gender, height, and initial P/F ratio were collected. Continuous ventilator data were collected using a laptop connected to the ventilator and using Hamilton DataLogger software (Hamilton, v5.0, 2011) to obtain pressure, flow, and volume measurements. Additionally, the DataLogger software allowed collection of ventilator mode and ventilator settings based on mode (i.e.: set tidal, respiratory rate, positive end-expiratory pressure (PEEP), and fraction inspired oxygen (FiO_2_)). Data were collected until extubation or for up to seven days per patient.

### Parameter estimation methodology

The damage-informed lung ventilator model is a complex model and we estimate its parameters for mouse and human clinical ventilator data. In clinical situations, the patient data are variable and often nonstationary because of interventions, patient-ventilator interactions, changes in health, etc., leading to complex parameter estimation issues. Moreover, the model we develop here is not likely to be structurally identifiable.^48–50^ However, formally computing identifiability properties here is subtle because many parameters in the model functionally affect only part of the breath. This feature helps facilitate the convergence of parameter estimates and potentially leads to the uniqueness of those estimates, although because the DILV model is neither linear nor convex, there is no guarantee of unique global optima and no way of guaranteeing that the optimal solution we compute is a, or the, global optimum. Nevertheless, this feature—parameters being active at different times during a breath—also makes formal structural identifiability calculations complex to compute. These complexities force us to address four issues, (1) computational estimation methodology, (2) management of parameter identifiability issues and parameter selection methods, (3) uncertainty quantification, and (4) estimation evaluation methodology.

#### Computational estimation methodology

Our needs require an estimation methodology that allows us to estimate states and parameters of the model effectively and the respective uncertainties in the estimated parameters. While stochastic methods, e.g., Markov Chain Monte Carlo (MCMC)^51^, might guarantee to find global minima and quantifying uncertainty in the estimated parameters values, they are generally quite slow. On the other hand, deterministic methods, e.g., Nelder-Mead optimization^52^, are substantially faster and by choosing many initial conditions, we are still able to quantify the uncertainty of a solution. In particular, here we infer parameters with a standard class of deterministic, multivariate, constrained nonlinear optimization methods, interior-point methods,^53,54^ a choice that is not critical among constrained, nonlinear optimization algorithms. As such, we focus on a smoothing task that employs deterministic nonlinear optimization methods that work well with careful parameter selection and constraints and can be used to quantify uncertainty.

#### Management of parameter identifiability and parameter selection methods

Irrespective of the estimation methodology, identifiability failure—non-uniqueness of non-convergence of solutions—can occur. In particular, the DILV model is likely not identifiable. To mitigate this problem, we use three different approaches to minimize impacts of identifiability failures while estimating parameters for a given waveform data. First, the model was constructed such that each parameter in the model contributes to the specific feature in the volume and pressure curves, allowing the parameter to be estimated relative to the specific, time-limited feature by definition. Second, we constrain the ranges of all parameters to lie within physically possible values. And third, we do not estimate every parameter in all circumstances but rather limit parameters estimated to those relevant for a given setting and fix many low-impact, low-sensitivity parameters.^55–57^

In more detail, the DILV model includes two state variables, volume and pressure with one overlapping parameter, the frequency of the breath (*θ*). The volume model has six parameters (*a_1_*, *b_1_*, *ϕ_1_*, *β_1_*, *β_2_*, *A_v_*). The pressure model has fourteen parameters (*a_2_*, *b_2_*, *ϕ_2_*, *a_3_*, *b_3_*, *ϕ_3_*, *β_3_*, *β_4_*, *β_5_*, *β_6_*, *A_p1_*, *A_p2_*, *A_p3_*, *A_p4_*). Many of these parameters can effectively be uniquely estimated because they operate on a particular part of the waveform, e.g., parameters that control the gradient of the rising signal (*β_1_*, *β_3_*) and falling signal (*β_2_*, *β_4_*) and amplitudes of the waveforms (*A_v_*, *A_p1_*). Nevertheless, there are parameters that are not necessarily uniquely estimable, e.g., parameters that control feature A1 (*a_3_*, *b_3_*, *ϕ_3_*, *A_p3_*) and features B1 and B2 (*β_5_*, *β_6_*, *A_p2_*).

#### Uncertainty quantification

Because we use deterministic optimization methods whose final solution depends on the initialization, we quantify uncertainty by randomly sampling a set optimization initialization for the parameters we estimate from a uniform distribution within a bounded interval (upper and lower bounds) centered around initial values.^58,59^ The boundaries of the intervals were chosen to exclude parameter variation that was unrealistic. The upper and lower boundaries of the intervals were chosen by computing parameters that provide a qualitative agreement between the model and the measured response. The optimal parameter estimate is then represented as a probability density in a similar way as is created using MCMC, allowing us to understand how informative, unique, and uncertain a given parameter solution set is. Additionally, we have uncertainty for individual breaths—we estimate every breath many times computing an uncertainty in by-breath parameter estimation—and uncertainty due to variation in many breaths over time. This allows us to both resolve and quantify single-breath features, and how those features vary over time, for different breaths, and even between individuals.

#### Estimation evaluation methodology

The output of this computational method is a *distribution* of optimal solutions. Through this distribution we understand the robustness of the solution and the uncertainty of the solution. If the distribution of solutions has multiple modes with similar error then we can conclude that there are multiple plausible solutions. Similarly, if the distribution of solutions is narrow or wide with similar errors, we can conclude that the model either does or does not depend highly on a given parameter. And finally, it is the distribution of parameter solutions that *define the phenotype computed by the model* in the sense that the distribution of parameters explains the by-breath characterization of the patient. We verify a model’s ability to represent data by computing the mean squared error between the model computed with parameter values taken as the medians of the optimally computed solution and the data. There is uncertainty in these MSE values too, and if one model has a lower MSE value than another with non-overlapping uncertainty in MSE, we conclude that the model with the lower median MSE more accurately represents the data.

## RESULTS

### Damage-informed lung ventilator model quantitative verification for experimental mouse model ventilator data

To demonstrate the effectiveness of the damage-informed lung-ventilator (DILV) model, we quantitatively verify the model by estimating parameters for data sets corresponding to different phenotypes—e.g., injured versus healthy. We then show differences in the estimated parameter values reflect different phenotypic states in a manner that is consistent with observed changes in lungs for mice in the absence and presence of ventilator dyssynchrony (VD) using data sets that belong to PVC and VCV modes of ventilation to verify volume and pressure models, respectively.

#### In PCV, volume model outcome agrees with the experimental outcome and with the single-compartment model

Figure 5, panel 1 shows two different pressure-controlled breaths recorded from the first mice in a healthy condition (green) and after a lung injury induced by injurious lavage and mechanical ventilation (orange). The pressure-volume loops suggest a reduction in lung compliance and an increase in hysteresis that is characteristic of lung injury. The DILV model estimated states (dashed-dot lines) show the same trends. The estimated parameter values and the respective uncertainty for individual breaths are shown in Table 2 with bold indicating physiological relevance.

**Table 2:** Estimated model parameters obtained from the optimization scheme for the results shown in Fig. 5 (Panel 1) and Fig. 6 (human patent 1) correspond to the mouse and human data, respectively. The error values were determined using the standard error of the mean for individual breaths. To quantify uncertainty in parameter estimates, each individual breath was estimated 1000 times. The parameters that are correlated with a known measure of lung physiology are in bold. Here, Cs and Cd are lung compliance values extracted by fitting the single-compartment model to data (Cs) and from the damaged-informed lung ventilator model (Cd) = *A_v_*/*A_p1_*.

| Parameters | Fig. 5 (Panel 1) |  | Fig. 6 (Patient 1) |  |
| --- | --- | --- | --- | --- |
|  | Healthy | Injured | Beginning | Ending |
| $\theta$ | 0.66 ± 0.000 | 0.66 ± 0.000 | 0.34 ± 0.000 | 0.35 ± 0.000 |
| $a_1$ | 9.67 ± 0.385 | 15.41 ± 0.397 | 7.51 ± 0.258 | 7.57 ± 0.282 |
| $b_1$ | 0.59 ± 0.006 | 0.53 ± 0.006 | 0.66 ± 0.000 | 0.63 ± 0.000 |
| $\Phi_1$ | 0.09 ± 0.007 | -0.01 ± 0.007 | 0.42 ± 0.003 | 0.51 ± 0.002 |
| $\beta_1$ | **112.97 ± 9.733** | **458.95 ± 8.179** | **10.96 ± 0.223** | **12.53 ± 0.240** |
| $\beta_2$ | **28.35 ± 10.490** | **10.60 ± 10.590** | **12.42 ± 0.051** | **12.64 ± 0.039** |
| $A_v$ | **1.03 ± 0.006** | **0.70 ± 0.003** | **402.61 ± 0.368** | **337.08 ± 0.531** |
| $a_2$ | 52.06 ± 1.440 | 32.11 ± 1.514 | 27.39 ± 0.118 | 17.73 ± 0.163 |
| $b_2$ | 0.80 ± 0.002 | 0.66 ± 0.001 | 0.67 ± 0.000 | 0.66 ± 0.000 |
| $\Phi_2$ | 0.34 ± 0.002 | 0.09 ± 0.002 | 0.35 ± 0.002 | 0.32 ± 0.001 |
| $a_3$ | **2.55 ± 0.334** | **5.11 ± 0.205** | **7.71 ± 0.183** | **18.37 ± 0.146** |
| $b_3$ | **1.00 ± 0.002** | **0.53 ± 0.003** | **0.91 ± 0.000** | **0.90 ± 0.000** |
| $\Phi_3$ | 0.00 ± 0.002 | 0.00 ± 0.005 | 0.15 ± 0.001 | 0.14 ± 0.001 |
| $\beta_3$ | **50.00 ± 0.388** | **100.84 ± 0.262** | **7.25 ± 0.157** | **12.88 ± 0.147** |
| $\beta_4$ | 18.54 ± 0.239 | 11.93 ± 0.193 | 1.95 ± 0.121 | 2.52 ± 0.064 |
| $\beta_5$ | - | - | 1.00 ± 0.000 | 1.00 ± 0.000 |
| $\beta_6$ | - | - | 1.13 ± 0.001 | 1.08 ± 0.005 |
| $A_{p1}$ | **35.53 ± 0.018** | **35.02 ± 0.017** | **16.58 ± 0.018** | **11.64 ± 0.008** |
| $A_{p2}$ | - | - | 3.80 ± 0.011 | 2.48 ± 0.022 |
| $A_{p3}$ | **14.00 ± 0.081** | **9.60 ± 0.077** | **3.90 ± 0.022** | **4.10 ± 0.016** |
| $A_{p4}$ | 0.06 ± 0.035 | 0.00 ± 0.021 | 19.98 ± 0.011 | 17.91 ± 0.004 |
| $C_s$ | 0.030 | 0.019 | 26.80 | 31.40 |
| $C_d$ | 0.029 | 0.020 | 24.28 | 28.96 |

**Figure 5:**
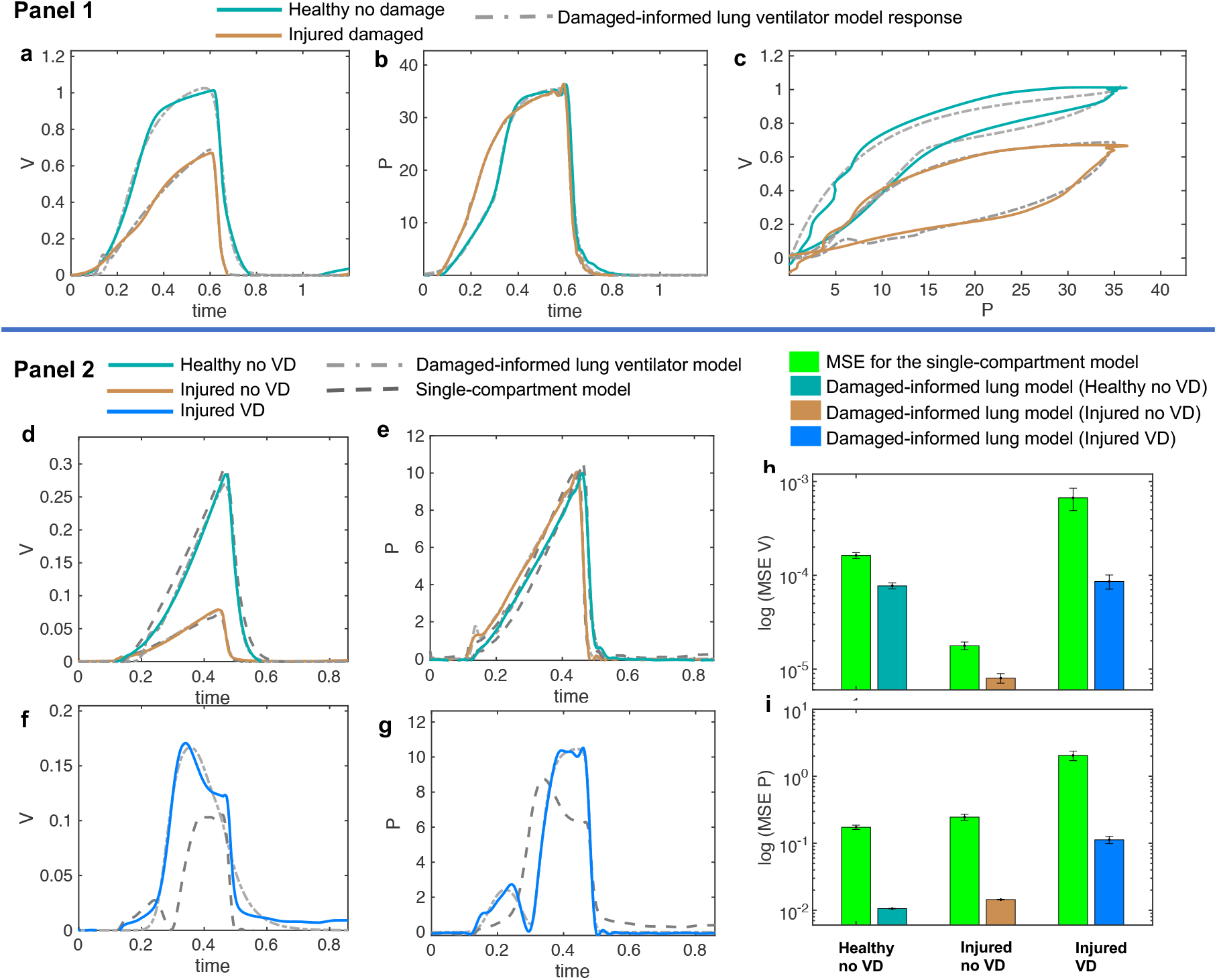
Volume and pressure models’ responses closely agree with the experimental data from the representative mice in healthy and injured condition. **(a-c)** In Panel 1, the measured breaths of first mouse are shown in healthy and injured conditions (solid lines) while the damaged-informed lung ventilator (DILV) model response is shown in dashed lines. Equations 1-18 were used to generate the best-fit model response using estimated mean parameter values shown in Table 1, respectively. In Panel 2, **(d-g)** three representative breaths of mouse two is shown here, which are corresponding to three different lung conditions (solid lines) while the damaged-informed lung ventilator model response is shown in the dashed-dot lines. For these breaths, histograms of the initial guesses and estimated parameters are shown in Supplementary Fig. S5a. The response of the single-compartment model is shown in dashed-dashed lines. **(f-i)** The cumulative values of mean squares errors (MSE) of the two models are compared here. These values corresponded to sixty breaths for healthy no ventilator dyssynchrony (VD) and injured no VD cases while ten breaths for injured VD case. For each case and breath, the estimated parameter values and MSE are shown in the Supplementary Fig. S6b-d. All the data shown here was collected in PCV (see Methods).

In the volume model, the injured lung showed a lower inspiratory flow rate, quantified by an increase in *β_1_*, and a faster expiration, quantified by a reduction in both *β_2_* and *A_v_* than the healthy lung model estimates. This might suggest a reduction in lung compliance, and the associated decrease in the expiratory time constant (See Table 1). Physiologic interpretation of the pressure model is limited because of the use of PCV. In this case, the pressure signal is prescribed by the ventilator and the observed differences between the healthy and injured lungs are a result of the ventilator control system algorithms. Hence, the respective changes in the parameters’ values, such as an increase in parameters *a_3_* and *β_3_* correspond to the changes in the ventilator settings and not in the respiratory mechanics. These results make an important point: it is essential to see the relative change in the parameters that control these features and to synthesize the model-based inference in a holistic fashion, instead of focusing on any one parameter or feature in isolation, given lung mechanics depends on both pressure and volume signal mutually.

To analyze changes in lung mechanics in a manner that accounts for ventilator settings, we define the lung compliance *Cd* = *A_v_*/*A_p1_* (Table 2), which is the ratio of volume and pressure model amplitudes. As expected, *Cd* decreases with injury. Furthermore, *Cd* shows a strong correlation with compliance (*Cs*) calculated with the single-compartment model^34^ (Table 2).

#### DILV model accurately captures mouse model data with ventilator dyssynchrony while the single-compartment model is unable to capture this variability

In section above, we demonstrate that the DILV model can estimate single breath accurately but we did not verify that the model has enough flexibility to account for the large variations in the waveform data. The DILV model can estimate a large number of breaths reliably while maintaining unique solutions of parameters values for each breath. To show these characteristics, we estimate a large number of breaths for the second mouse in three different lung conditions: 1) healthy breaths without VD, 2) injured breaths without VD, and 3) injured breaths with VD (See Methods). In the 1^st^ and 2^nd^ cases, we selected sixty breaths in a sequence from the random location in the data. In the 3^rd^ we manually selected ten breaths that had VD from a large data set containing breaths with and without respiratory effort. A representative breath for each case is shown in Fig. 5, panel 2 along with the DILV model response at the optimum parameter values (Supplementary Table S1). Histograms of the initial guesses and estimated parameters are shown in the Supplementary Fig. S5a. We observed minor variability in most of the estimated parameters values for individual breaths, suggesting that each feature is modularly controlled by the respective parameter. We do observe high variability in some parameters (*a_1_*, *β_1_*, *a_3_*, *β_3_*) due to the low sensitivity of the model for those parameters (Supplementary Fig. S5a).

To demonstrate what is gained by the DILV model we compare it to single-compartment model^34^ estimated resistance and compliance to estimate the same breaths for each case (Fig. 5, panel 2). The single-compartment model has substantially larger estimated mean squares errors (MSE) and these errors increase as the mouse lung condition worsens (Fig. 5h,i, Supplementary Fig. S5b-d). Consequently, in terms of lung compliance values, the two models’ outcomes closely agree in the healthy lung case but then diverge for the injured lung, and especially during VD (Supplementary Fig. S5e-g). These discrepancies have their root in the limited ability of the single-compartment model to estimate respiratory efforts (Fig. 5f,g), errors that are quantified by the MSE between model estimates and data (Supplementary Fig. S5b-d). Note that the compliance value that we calculate includes the effects of muscle effort – it’s not a true pulmonary system compliance. It’s also different from the non-VD breaths in that same injured mouse.

#### In VCV, pressure model outcome agrees with the experimental outcome

We verified the DILV model using data sets that were collected during PCV, where the estimated volume model parameters reflected the lung dynamics since the volume was the free variable. We now consider data set that was collected in VCV mode where the estimated pressure model parameter should reflect the lung condition. For that, we preform third mouse verification including variations in PEEP during low tidal volume ventilation (VCV). The PV loops show reduced compliance with injury for both PEEP settings. The full PEEP-varied results are shown in the Supplementary Fig. S6. The pressure model indicates a reduction in compliance in the injured lung as quantified by lower values of parameters *a_3_, b_3_* and *β_3_*, and elevated estimates of in *A_p1_* (Table 1, Supplementary Fig. S6, Supplementary Table S1). In contrast to the results shown in Fig. 5, differences in parameter estimates between healthy and injured lungs in the pressure model were much larger compared to those estimated differences in the volume model. This is expected since the tidal volumes were approximately equal during VCV, and the reduction in compliance is reflected in increased pressure. This effect can be inferred by analyzing the *A_v_*/*A_p1_* ratio showing a reduction in the injured cases at both the PEEP settings (Supplementary Table S1). We also note that the healthy lung becomes stiffer at PEEP = 12 cmH_2_O due to strain stiffening. In contrast, the injured lung becomes more compliant at high PEEP, presumably due to recruitment.

### Damage-informed lung ventilator model quantitative verification for intensive care unit (ICU) patient-ventilator data

The DILV model is intended to be used with both laboratory data and clinical ventilator data where standard models, such as the single-compartment model, cannot recapitulate all of the potentially relevant waveform features, particularly patient-ventilator interactions and pathophysiological lung problems. To show the clinical applicability of the DILV model we consider waveform data of two ICU patients, the first—patient 1—includes waveform data without VD and the second—patient 2—has waveform data with VD. For each case, we estimate each individual breath 1000 times to quantify uncertainty in parameter estimates for each of 263 breaths to quantify uncertainty in parameter estimates across many breaths. These data were recorded near extubation when ARDS had nearly resolved. For both patients, the ventilator was operating in patient-triggered mode, a ventilator mode that is not possible in our mouse ventilators but is commonly used ventilator mode in the ICU.

#### Patient 1 ICU data-driven verification in the absence of VD

For patient 1, of the thousands of breaths available, we selected a sequence of 130 breaths without VD at a random location and performed parameters estimation breath by breath. The sequence of breaths starts at PEEP = 20 cmH_2_O and switches to PEEP = 18 cmH_2_O at breath number 68. Fig. 6a-c shows two representative breaths along with the DILV model response at the optimum parameter values (Table 2). Histograms of the initial guesses and estimated parameters are shown in the Supplementary Fig. S7a for the respective breaths. For all the model parameters, we observed unimodal estimated parameter distributions with low variance, suggesting that each parameter controls a specific feature of the waveform. We also estimated each breath using the single-compartment model and substantially higher MSE compared to the DILV model (Fig. 6g,h). Note that in Fig. 6g-j, an MSE ratio less than one indicates that the DILV model produced waveforms that were more similar to the measured data.

**Figure 6:**
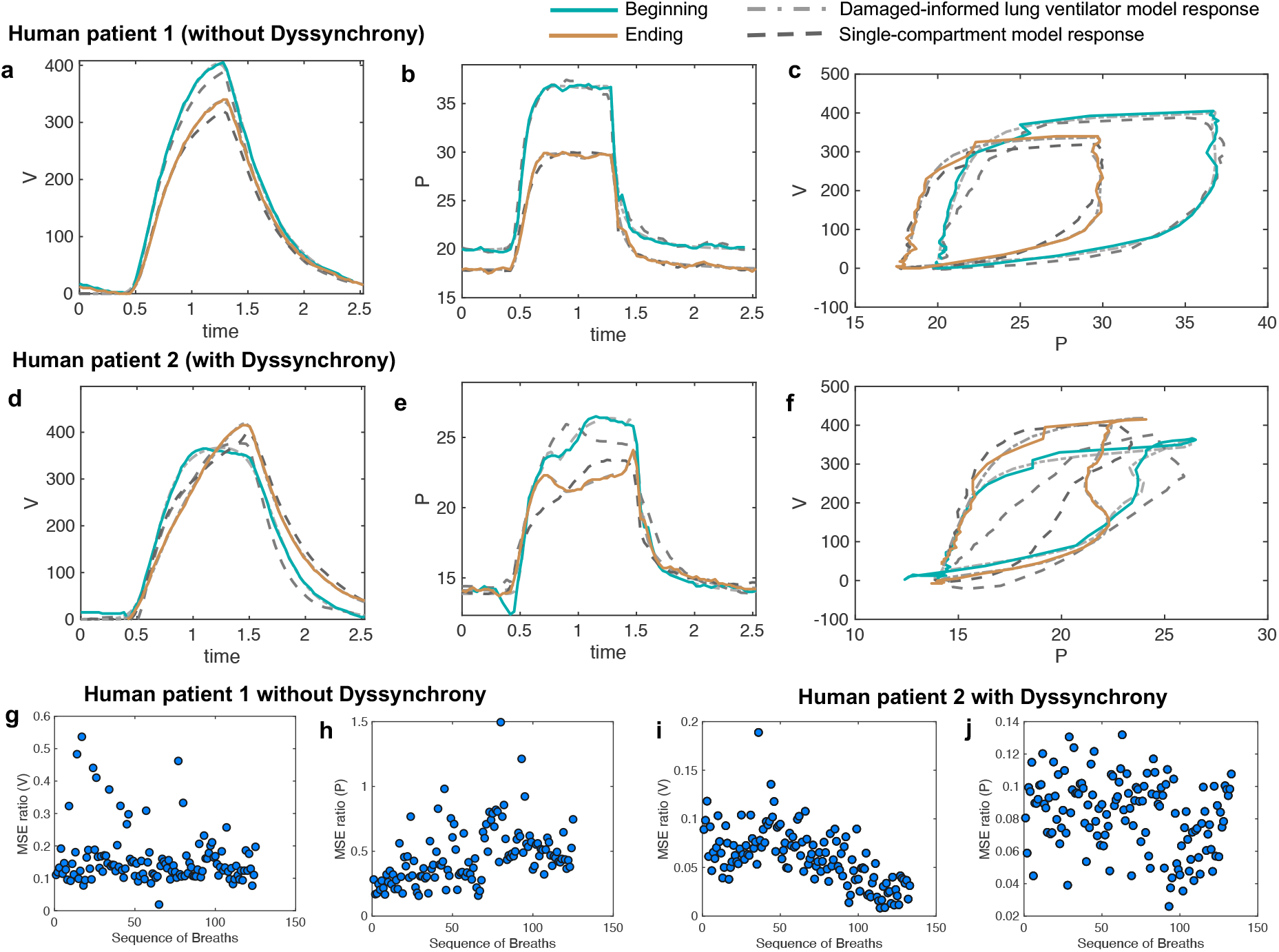
Damaged-informed lung ventilator model can accurately follow breaths of the ICU patients with ARDS without and with dyssynchrony. Measured representative breaths of **(a-c)** patient 1 without dyssynchrony and **(d-f)** patient 2 with dyssynchrony are shown in solid lines, while the DILV model inferred response is shown in the dashed-dot lines. The response of the single-compartment model is shown in dashed-dashed lines. Histograms of the initial guesses and estimated parameters are shown in the Supplementary Figs. S7a and 8a. For all the breaths in each case, the corresponding ratio of mean squares errors (MSE) between the DILV model and the single-compartment model is shown in **(g-h)** and **(i-j)**, respectively, while histograms are shown in Supplementary Figs. 9 and 10. The response of the DILV model was determined using Eqns. 1-18 to generate the best-fit model response using estimated mean parameter values shown in Table 1, and Supplementary Table 1. All the data shown here were collected in human-triggered mode (see Methods).

PV loops for these cases (Fig. 6c) suggest that lung compliance is increased by the prescribed change in PEEP. We found that the model-estimated parameters indicate an increase in compliance (the *A_v_*/*A_p1_* ratio) with a reduction in PEEP (Table 2). The increased compliance at lower PEEP agrees with the single-compartment model^34^ response (Supplementary Fig. S8b,c) and also the patient outcome (successful extubation). Moreover, a significant increase in parameters *a_3_,* and *β_3_* suggested the same (Table 2). The prescribed reduction in PEEP was reflected in a reduction in *A_p4_*.

#### Patient 2 ICU data-driven verification in the presence of VD

In the cases where some patient effort is present, e.g., VD, additional model parameters must be used to quantify and understand the patient-ventilator interaction. To show this, from patient 2’s data we randomly selected 133 breaths from the thousands of breaths observed. These breaths were selected from a several hours long time window where most of the breaths showed mild to severe VD. Two representative breaths for this case are shown in Fig. 6d-f along with the DILV model’s estimate of these breaths at the optimum parameter values (Supplementary Table S1). Histograms of the initial guesses and estimated parameters for these breaths are shown in the Supplementary Fig. S8a.

PV loops for these breaths (Fig. 6f) suggest that lung compliance is increased prior to extubation compared to earlier in the ICU course. This qualitative observation is supported by the increased *A_v_*/*A_p1_* ratio (Supplementary Table 1) that is a measure of compliance. However, it is important to note that this compliance includes the additional effects of respiratory effort. To further demonstrate the importance of the DILV model, we compared the DILV model’s estimate of the breaths with the single-compartment model^34^ estimates for each breath. The single-compartment model estimates of the breaths have substantially higher MSE values for patient 2 compared to patient 1 (Fig. 6g-j) due to the presence of respiratory efforts. Accordingly, we expect large errors in the compliance values estimated with the single-compartment model (Supplementary Fig S8b,c). These results agree with the fact that the DILV model can estimate the volume and pressure waveform data more accurately, especially when the waveforms have high variability, as in the case of patient 2 with VD.

The mouse data verification showed that the DILV model is able to estimate most of the parameters for individual breaths. It is also important to quantify uncertainty across many breaths and in different patients. Analyzing the variability in the estimated parameters over several breaths might capture the lung condition’s heterogeneous nature, including many potentially different and differently damaging breaths that show VD. In the ICU, there are no controlled experiments, and patients simultaneously experience many types and severities of VD, different interventions, heterogeneous injurious insults, etc. We expect to see minor variability in parameter estimates when many breaths of a patient are approximately the same compared to large variability in the parameter estimates when breaths are heterogeneous. For example, we observed a low variability in all the volume model parameters over several breaths with the exception of *A_v_* for patient 1, indicating that the volume waveforms’ characteristic shape remains the same at different time points except for variations in tidal volume. This contrasts with what we observed for patient 2 where VD drove heterogeneity and substantial deviations from more normal breaths (Supplementary Figs. S9, S10). More importantly, parameters associated with patient-ventilator dyssynchrony can be used to infer lung condition. For example, for patient 2, we observed that estimated distributions of *β_5_*, *β_6_*, *A_p2_* parameters across several breaths have higher values than patient 1 and indicating the VD known to be present in patient 2 (Supplementary Figs. S9, S10).

Overall, these results suggest that the DILV model can reproduce a wide variety of waveform data and is capable of extracting hypothesis-driven, clinically-relevant information from the waveforms that might facilitate a systematic interpretation of the dynamics of the injured lung.

## DISCUSSION

We developed a damage-informed lung ventilator model that represents pressure and volume time-series data by reconstructing the waveforms from hypothesis-driven modular subcomponents. We demonstrated the efficacy of the model using simulations for qualitative validation and by estimating mouse and human data for quantitative verification. The model accurately estimated volume and pressure waveforms data and consistently distinguished healthy from injured lungs based on parameter estimation. Furthermore, we directly incorporate clinical and physiologic knowledge regarding important and observable features into the model that might be associated with the lung pathophysiology—the subcomponents of the model represent hypothesis-driven deviations from normal breaths. We also analytically define lung compliance in terms of model parameters and demonstrate changes in compliance values that agree with the known lung condition.

To demonstrate what is gained with this novel approach, we present a comprehensive comparison between the DILV model and the single-compartment model for a wide range of ventilator waveforms related to different lung conditions and patient-ventilator interactions. We include pressure- and volume-controlled ventilation in healthy and lung-injured mice and in humans in the absence and presence of respiratory effort. Through this comparison, we establish that the DILV model can reproduce all the features present in the waveform data and report lung compliance values that agree with lung condition (Fig. 5, 6 and Supplementary Fig. S5, S7-S10). This is primarily possible because of our unique waveform-based approach that enabled the model to have enough flexibility. At the same time, the model is limited using prior knowledge so as not to have the capability to estimate every possible variation in PV waveforms, but rather is constrained to estimate the features of the ventilator data that are the most clinically impactful. This approach lives between a machine learning approach, where the model is flexible enough to estimate every feature and must then discern which features are important through regularization to prevent overfitting, and the fully mechanistic lung modeling approach where the observed physiology must emerge from the proposed lung mechanics. It is possible that taking this middle path will help advance all approaches.

The most direct application of the DILV modeling approach is to quantify the qualitative physiological interpretation of pressure and volume data. An experienced clinician or physiologist can infer the status of a patient, the safety of ongoing ventilation, the presence of ventilator dyssynchrony, and other important details from visual inspection. However, we currently do not yet have methods to identify all of these characteristics in ventilator data quantitatively. The entire waveform may be utilized and this provides a rich repository of data that is challenging and time-consuming to use for diagnosis and treatment. In contrast, summarizing the waveform data in scalar values for resistance and compliance may cast aside important details such as identifying dyssynchrony. Our approach may offer a methodology for condensing the pressure-volume data to track changes in injury severity over time, and estimate injury phenotypes (Supplementary Figs. S9, S10). A similar phenotype study on a large dataset could be used to categorize and understand lung injury, serve as outcome measures for interventions, and describe the impacts of VILI and dyssynchrony,^38^ and VILI. ^1–7,60,61^

Lung injury diagnosis and decision-making are based in part on interpretation of the pressure, volume, and flow waveforms. However, different pathophysiologic mechanisms can lead to the same observed waveform features. For example, increased driving pressure could be a result of derecruitment (alveolar collapse) or alveolar flooding. ^62,63^ In other words, the human-based inference using limited waveform data can be ill-posed. The DILV modeling and parameter estimation approach could allow to estimate a large number of breaths efficiently with unique solutions (Supplementary Figs. S9, S10). We could therefore use the DILV model to estimate over many similar but varied breaths, and might be to better triangulate the most probable pathophysiologic drivers because the primary driver of damage will likely be present, and significant, despite inter-breath variations. At the same time, more extraneous details will not be consistently expressed in every breath.

Note that in the DILV model, an explicit coupling between pressure and volume signals is absent. We have intentionally taken this approach to preserve flexibility so that we can accurately recapitulate a wide variety of clinically and experimentally observed features in the pressure and volume signals, including the effects of ventilator dyssynchrony (Figs. 5f,g, and 6d-f). Such flexibility in the model outcome is not always possible with rigid coupling between pressure and volume data, as we have shown by comparing the DILV model response with the single-compartment model. This is not to say that pressure and volume are totally independent in the DILV model because we utilize the same respiratory rate for both. In addition, we show that the ratio of a volume and pressure model parameter (*A_v_* and *A_p1_*) describes lung compliance. In future studies, we will link additional specific components of the pressure and volume waveforms through physiologically relevant parameters such as nonlinear lung elastance or inspiratory and expiratory flow resistance. Alternatively, compartment-based model could be used to incorporate the physiologic coupling between pressure and volume data by utilizing the outputs from the model presented here as inputs for compartment models. Moreover, if the DILV model is used to preprocess the data before analysis using a compartment model, it is possible to formulate the problem entirely of ordinary differential equations, opening up a range of more efficient inference machinery. ^53–55^

Finally, our work here has several notable limitations. *First*, the DILV model quantitative verification was conducted using healthy and severely lung-injured mice as well as a limited human data set. This is sufficient for proof in principle that the model can capture physiologic differences. However, establishing that the model can accurately differentiate more specifically defined phenotypes will require verification on much larger laboratory and clinical populations. *Second*, the human data in this study were collected using a specific ventilator (Hamilton G5) operated in a certain mode. For wider applicability of the model, additional data verification is required across different ventilators and modes. *Third*, we relied on the expert knowledge of a single critical care physician to determine the clinically important characteristics of the pressure and volume waveforms and it is likely that differing opinions will exist among intensivists. Collecting and synthesizing such information will require a different qualitative study. Moreover, there may be differing opinions regarding what should and should not be included in the model. This does not negate our methodology or the DILV model. Instead, it suggests future work is necessary to understand better and verify clinically important features. Alternatively, we may instead seek to link model features to patient outcomes, thus establishing the important characteristics of the model by linking those parameters to outcomes.

In summary, we developed a physiologically anchored and data-driven lung ventilator model that can reproduce the important features pressures and volumes during mechanical ventilation. The performance of the model was verified with experimental and clinical data in healthy and injured lungs to demonstrate model efficacy in robustly estimating interpretable parameters. This methodology represents a departure from many lung modeling efforts, and suggests future directions of work that can provide another pathway for better understanding lung function during mechanical ventilation and can potentially form a bridge between experimental physiology and clinical practice.

## Author Contribution

D.K.A., B.J.S., and D.J.A. conception and design of research; D.J.A and D.K.A. developed the mathematical model, D.K.A., B.J.S., and P.D.S. designed and conducted experiments; D.K.A., B.J.S., and D.J.A. analyzed data and evaluated the model; D.K.A., B.J.S., P.D.S., and D.J.A. interpreted results; D.K.A. prepared figures; D.K.A. wrote the original draft of the manuscript; D.K.A., B.J.S., P.D.S., and D.J.A. edited and revised the manuscript; D.K.A., B.J.S., P.D.S., and D.J.A. approved final version of manuscript.

