## Supplementary File for "A damaged-informed lung ventilator model for ventilator waveforms"

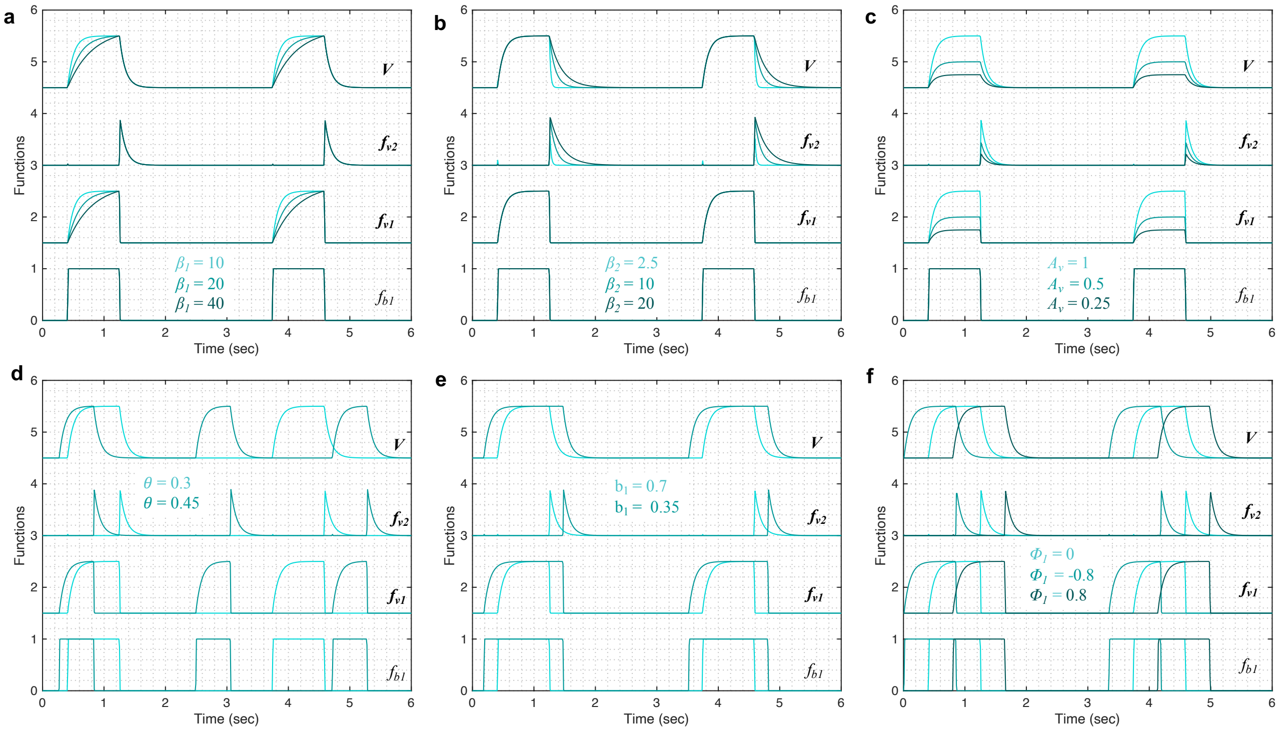


Figure S1: Showing the effect of parameter variations on the submodels that form the volume model for the cases shown in Fig. 3. The gradient of the rising and falling signals can be altered using the (a) $\beta_1$ and (b) $\beta_2$ parameters, respectively. Increased values of these parameters increase the transient time for the signal to reach the same volume level. (c) The amplitude of the waveform can be altered using the parameter $A_v$. (d) Changes in the respiratory frequency ($\theta$) change the period of the breath while (e) the I:E ratio (inspiratory to expiratory time ratio) can be modified using the $b_1$ parameter. The starting point of the inspiration can be modified using the parameter $\phi_1$. The output of the model ($V$) was calculated using Eqns. (1)-(5) while considering $\theta = 0.3$, $a_1 = 200$, $b_1 = 0.7$, $\phi_1 = 0$, $\beta_1 = 10$, $\beta_2 = 10$, $A_v = 1$. Y-axis was normalized to represent all the submodels in a sequential manner. Note that the model variability shown here is independent of the ventilator mode.

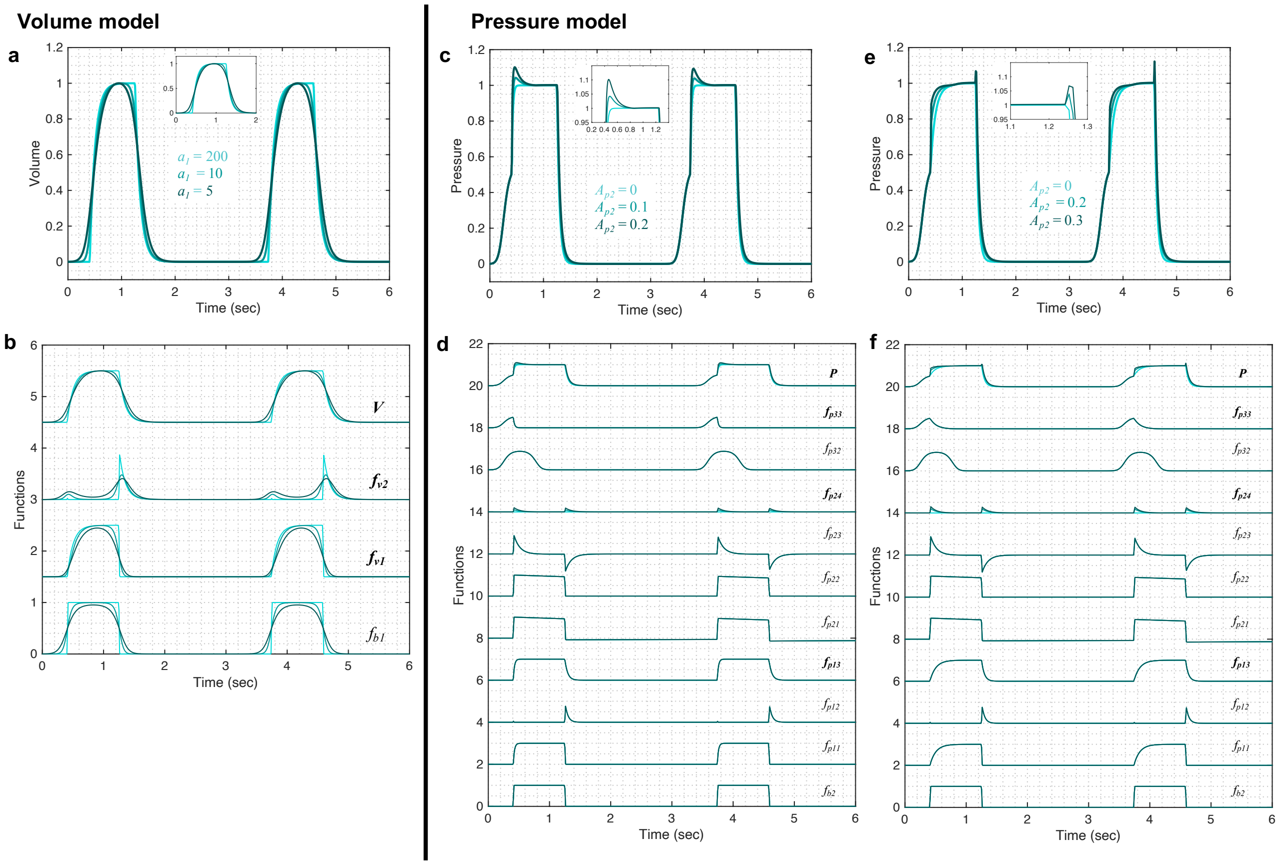


Figure S2: (a-b) Further variations in the volume waveform can be achieved by changing the (a) $a_1$ parameter while in the pressure waveform, control over B1 and B2 features via (c-f) $A_{p_2}$ parameter when (g) $\beta_3 = 5$, (h) $\beta_3 = 2.5$ (i) $\beta_3 = 10$. The output of the model ($V$) was calculated using Eqns. (1)-(5) while considering $\theta = 0.3$, $a_1 = 200$, $b_1 = 0.7$, $\phi_1 = 0$, $\beta_1 = 10$, $\beta_2 = 10$, $A_v = 1$. Equations (6)-(18) were used to simulate the response of the pressure model while considering $\theta = 0.3$, $a_2 = 200$, $b_2 = 0.7$, $\phi_2 = 0$, $a_3 = 10$, $b_3 = 0.9$, $\phi_3 =$ -0.6, $\beta_3 = \beta_4 = 5$, $\beta_5 = 1.001$, $\beta_6 = 1.1111$, $A_{p_1} = 1$, $A_{p_2} = 0$, $A_{p_3} = 0.5$, $A_{p_4} = 0$. Y-axis was normalized to represent all the submodels in a sequential manner. Note that the model variability shown here is independent of the ventilator mode.

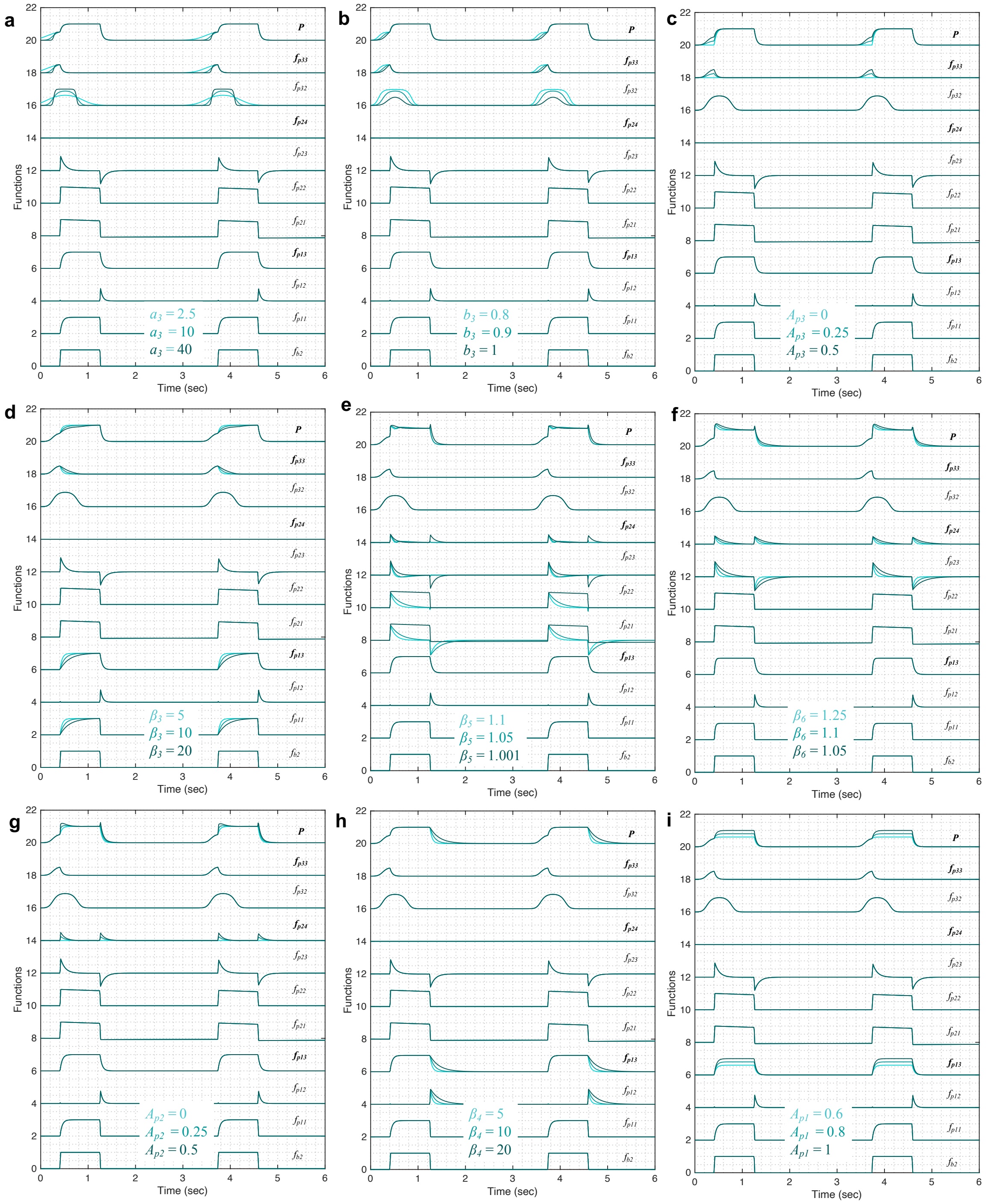


Figure S3: Showing the effect of parameter variations on the submodels that make up the pressure model for the cases shown in Fig. 4. The initial gradient of the pressure signal during inspiration at low volume (A1) is controlled by (a) the $a_3$ and (b) $b_3$ parameters while (c) the inflection point location can be altered using the $A_{p_3}$ parameter. (d) The gradient of the rising signal after the inflection point (A2), is controlled by the $\beta_3$ parameter. (e) The shapes of the peaks at the beginning (B1) and at the end (B2) of the plateau are regulated by the $\beta_5$ parameter when $A_{p_4} = 0.5$. (f) The sharpness of these peaks can be altered further by the $\beta_6$ parameter for a given shape while (g) the amplitude of the peaks is controlled by the $A_{p_2}$ parameter. (h) The gradient of the falling signal (C) during expiration can be modified by the $\beta_4$ parameter. (i) The amplitude of the waveform can be altered using the parameter $A_{p_1}$. Equations (6)-(18) were used to simulate the response of the pressure model while considering $\theta = 0.3$, $a_2 = 200$, $b_2 = 0.7$, $\phi_2 = 0$, $a_3 = 10$, $b_3 = 0.9$, $\phi_3 =$ -0.6, $\beta_3 = \beta_4 = 5$, $\beta_5 = 1.001$, $\beta_6 = 1.1111$, $A_{p_1} = 1$, $A_{p_2} = 0$, $A_{p_3} = 0.5$, $A_{p_4} = 0$. Y-axis was normalized to represent all the submodels in a sequential manner.

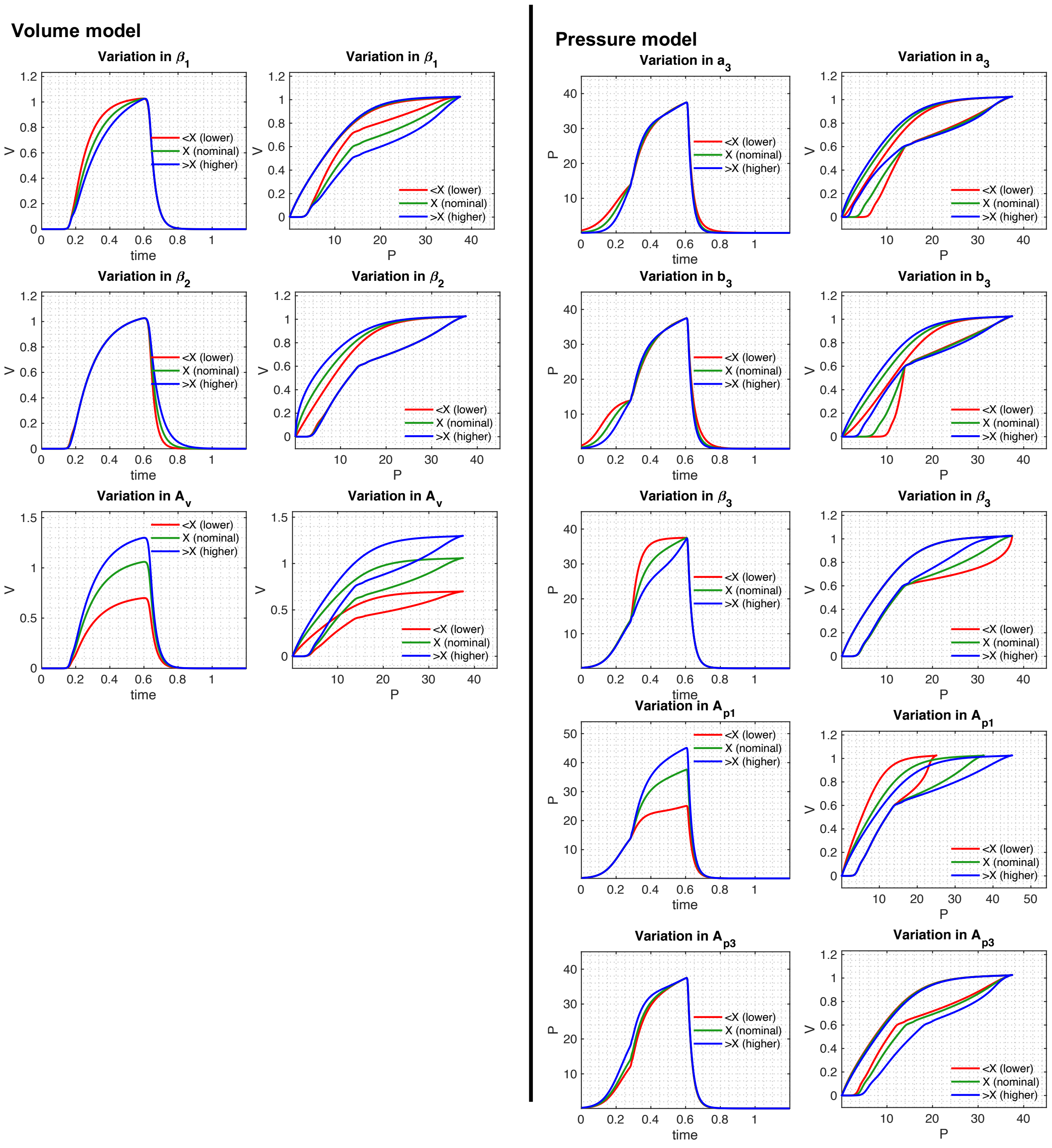


Figure S4: Three parameters in the volume model could have direct physiological meaning. These are $\beta_1$, $\beta_2$ and $A_v$. Five parameters in the pressure model could have direct physiological meaning. These are $a_3$, $b_3$, $\beta_3$, $A_{p_1}$ and $A_{p_3}$. Simulated response of the volume and pressure models are shown when the specific parameter was varied while considering the nominal parameter values shown in Table 1 for the mouse model PCV, healthy case (Fig. 6 Panel 1). Here, variable $X$ corresponds to the nominal value of the parameter while $< X$ and $> X$ represent smaller and larger values than the nominal value, respectively. Note that the model variability shown here is independent of ventilator mode. While analyzing one variable (volume or pressure), the other one (pressure and volume) is considered to be fixed.

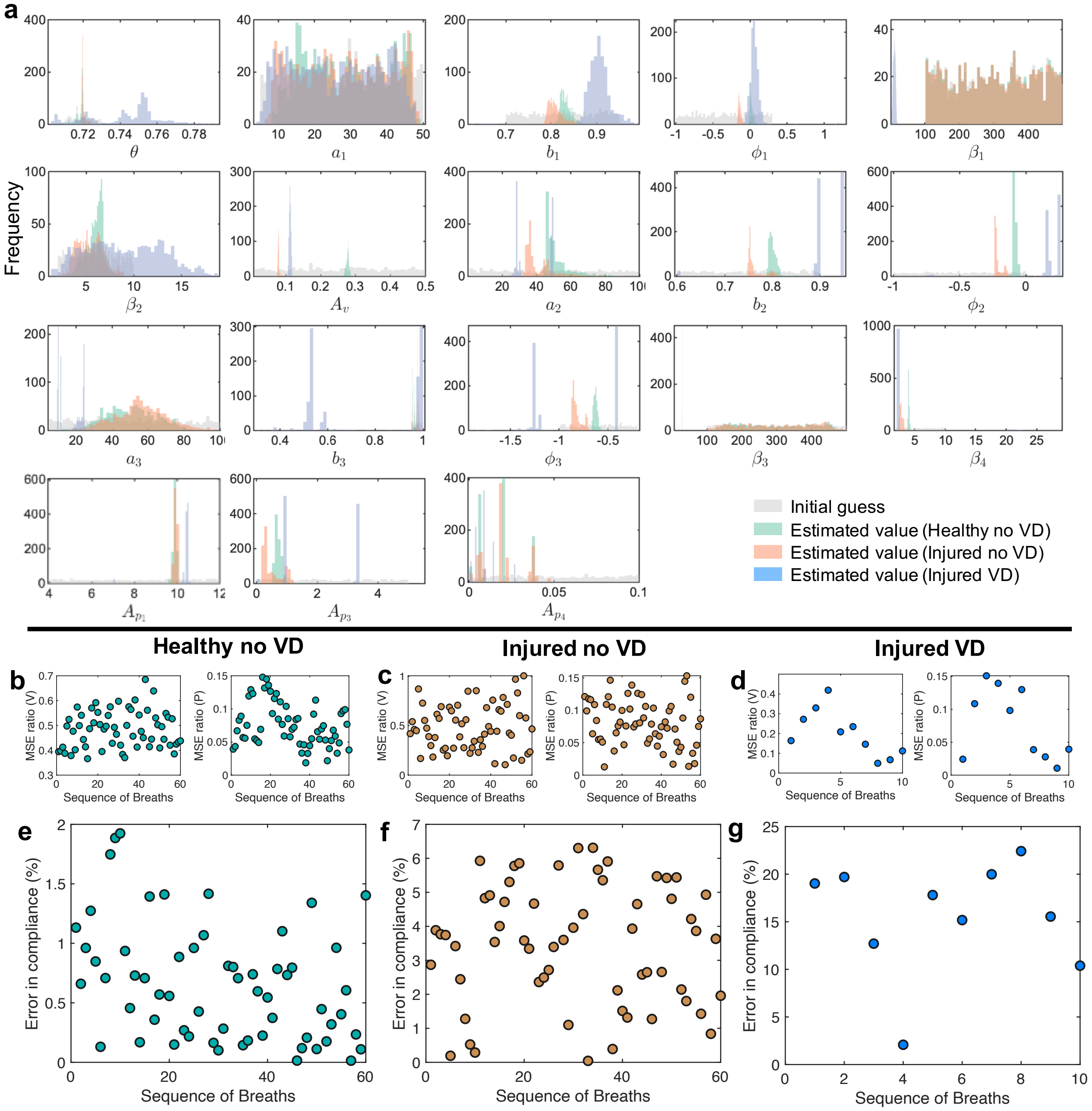


Figure S5: Histograms of the initial guesses and estimated parameters were obtained from 1000 samples that gave the lowest fitting error (see Methods).These results correspond to the breaths shown in Panel 2 of Fig. 5 in the main text. (b-g) The ratio of the mean squares errors between our model and the single-compartment model for 60 breaths each in healthy no VD and injured no VD conditions and 10 breaths in injured VD case. (e-g) Percentage error in the single-compartment lung compliance values with respect to our model. In the latter case, $A_v/A_{p_1}$ ratio was used to calculate the compliance using the estimated parameters values for each breath.

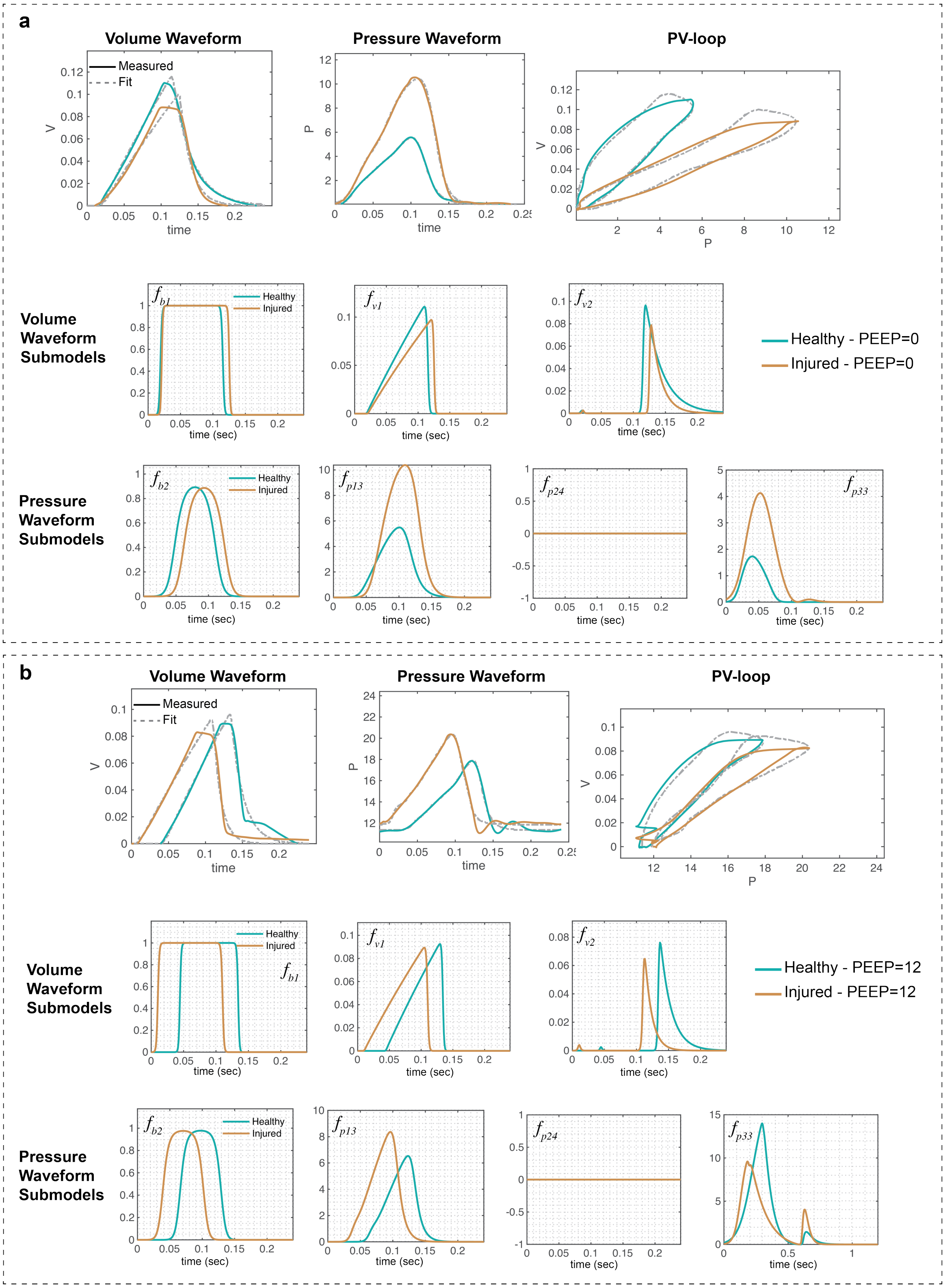


Figure S6: Experimental data from a representative mouse, under VCV, is compared with the model data in healthy and injured condition at (a) PEEP $= 0$ and (b) PEEP $= 12$. In each panel, in the first row the measured response is shown in solid lines while the model inferred response is shown in dashed lines. Changes in the volume and pressure submodels are shown in the second and third rows, respectively (in solid lines). The volume and pressure models shown in Eqns. (1)-(5) and (6)-(18) were used to generate the best-fit model response using estimated mean parameter values shown in Table S1, respectively. The respective uncertainties in the parameter values are shown in Table S1.

Table S1: Estimated model parameters obtained from the optimization scheme for the results shown in Fig. 5-panel 2, Supplementary Fig. S6, and Fig. 6-patient 2. The error values were determined using the standard error of the mean. The parameters that are correlated with a known measures of lung physiology are in bold. N =1000. Here, $Cs$ and $Cd$ are lung compliance values extracted by fitting the single-compartment model to data and from the damaged-informed lung ventilator model. In latter case, $A_v/A_{p_1}$ ratio was used to calculate the compliance using the parameters values shown in the table.

| | Fig. 5 (Panel 2) | | | Supplementary Fig. S6a | | Supplementary Fig. S6b | | Fig. 6 (Patient 2) | |
|---|---|---|---|---|---|---|---|---|---|
| Parameters | Healthy no VD | Injured no VD | Injured VD | Healthy (PEEP=0) | Injured (PEEP=0) | (PEEP=12) | (PEEP=12) | Beginning | Ending |
| $\theta$ | 0.72 ± 0.000 | 0.72 ± 0.000 | 0.74 ± 0.001 | 0.72 ± 0.000 | 0.72 ± 0.000 | 0.72 ± 0.000 | 0.72 ± 0.000 | 0.35 ± 0.000 | 0.31 ± 0.000 |
| $a_1$ | 19.81 ± 0.354 | 26.29 ± 0.380 | 13.34 ± 0.381 | 21.50 ± 0.696 | 27.72 ± 0.698 | 26.92 ± 0.697 | 22.27 ± 0.700 | 3.32 ± 0.394 | 8.23 ± 0.384 |
| $b_1$ | 0.83 ± 0.000 | 0.79 ± 0.001 | 0.95 ± 0.001 | 0.47 ± 0.002 | 0.39 ± 0.002 | 0.53 ± 0.002 | 0.43 ± 0.002 | 0.45 ± 0.002 | 0.55 ± 0.002 |
| $\Phi_1$ | 0.02 ± 0.001 | -0.15 ± 0.001 | 0.03 ± 0.003 | -0.03 ± 0.002 | 0.10 ± 0.002 | 0.46 ± 0.002 | -0.19 ± 0.002 | 0.81 ± 0.006 | 0.40 ± 0.003 |
| **$\beta_1$** | **483.08 ± 3.706** | **454.31 ± 3.708** | **1.15 ± 0.123** | **498.47 ± 0.881** | **406.49 ± 0.883** | **411.04 ± 0.881** | **438.79 ± 0.884** | **4.77 ± 0.343** | **26.51 ± 0.235** |
| **$\beta_2$** | **5.63 ± 0.030** | **3.21 ± 0.042** | **15.05 ± 0.108** | **27.30 ± 0.320** | **16.22 ± 0.335** | **17.91 ± 0.335** | **11.61 ± 0.350** | **8.03 ± 0.110** | **13.48 ± 0.100** |
| **$A_v$** | **0.28 ± 0.000** | **0.08 ± 0.000** | **0.17 ± 0.000** | **0.12 ± 0.000** | **0.10 ± 0.000** | **0.10 ± 0.000** | **0.09 ± 0.000** | **367.01 ± 1.245** | **419.65 ± 1.055** |
| $a_2$ | 46.10 ± 0.253 | 32.09 ± 0.288 | 49.25 ± 0.106 | 5.07 ± 0.836 | 5.00 ± 0.843 | 8.27 ± 0.836 | 8.14 ± 0.841 | 11.48 ± 0.059 | 11.82 ± 0.068 |
| $b_2$ | 0.85 ± 0.000 | 0.75 ± 0.001 | 0.93 ± 0.001 | 0.79 ± 0.004 | 0.79 ± 0.006 | 0.77 ± 0.002 | 0.77 ± 0.004 | 0.69 ± 0.000 | 0.52 ± 0.001 |
| $\Phi_2$ | 0.00 ± 0.001 | -0.23 ± 0.001 | 0.41 ± 0.005 | 0.24 ± 0.004 | 0.56 ± 0.009 | 0.65 ± 0.008 | 0.04 ± 0.010 | 0.79 ± 0.002 | 0.48 ± 0.002 |
| **$a_3$** | **14.93 ± 0.459** | **93.86 ± 0.458** | **17.59 ± 0.075** | **6.78 ± 0.037** | **2.96 ± 0.047** | **9.06 ± 0.005** | **9.99 ± 0.011** | **28.63 ± 0.178** | **28.53 ± 0.149** |
| **$b_3$** | **0.95 ± 0.000** | **1.00 ± 0.000** | **1.00 ± 0.007** | **0.88 ± 0.004** | **0.70 ± 0.007** | **0.88 ± 0.011** | **0.41 ± 0.003** | **0.97 ± 0.000** | **0.98 ± 0.001** |
| $\Phi_3$ | -0.47 ± 0.001 | -0.79 ± 0.002 | -0.46 ± 0.010 | -0.46 ± 0.003 | 0.00 ± 0.001 | 0.00 ± 0.003 | 0.00 ± 0.004 | -0.15 ± 0.012 | -0.05 ± 0.008 |
| **$\beta_3$** | **124.10 ± 3.283** | **489.33 ± 3.366** | **6.45 ± 0.115** | **43.28 ± 5.080** | **20.01 ± 5.499** | **145.40 ± 1.256** | **139.74 ± 0.785** | **7.29 ± 0.032** | **17.42 ± 0.091** |
| $\beta_4$ | 3.80 ± 0.009 | 2.35 ± 0.011 | 2.01 ± 0.108 | 12.78 ± 5.402 | 10.21 ± 10.497 | 10.00 ± 9.056 | 10.00 ± 9.735 | 2.28 ± 0.046 | 2.21 ± 0.029 |
| $\beta_5$ | - | - | - | - | - | - | - | 1.0017 ± 0.0001 | 1.0025 ± 0.0001 |
| $\beta_6$ | - | - | - | - | - | - | - | 1.1029 ± 0.0051 | 1.1032 ± 0.0040 |
| **$A_{p1}$** | **9.95 ± 0.009** | **10.01 ± 0.006** | **10.45 ± 0.019** | **5.59 ± 0.017** | **10.42 ± 0.109** | **6.85 ± 0.112** | **8.77 ± 0.082** | **11.88 ± 0.012** | **8.27 ± 0.005** |
| $A_{p2}$ | - | - | - | - | - | - | - | 3.45 ± 0.032 | 3.67 ± 0.018 |
| **$A_{p3}$** | **1.47 ± 0.007** | **0.24 ± 0.011** | **2.42 ± 0.025** | **1.73 ± 0.016** | **4.13 ± 0.070** | **1.28 ± 0.142** | **1.77 ± 0.112** | **7.96 ± 0.029** | **1.98 ± 0.024** |
| $A_{p4}$ | 0.00 ± 0.000 | 0.00 ± 0.000 | 0.00 ± 0.000 | 0.06 ± 0.001 | 0.05 ± 0.004 | 11.36 ± 0.154 | 11.89 ± 0.081 | 13.86 ± 0.007 | 14.12 ± 0.002 |
| $Cs$ | 0.0289 | 0.0079 | 0.0194 | 0.021 | 0.009 | 0.015 | 0.011 | 33.04 | 48.36 |
| $Cd$ | 0.0285 | 0.0081 | 0.0164 | 0.021 | 0.010 | 0.014 | 0.011 | 30.89 | 50.77 |

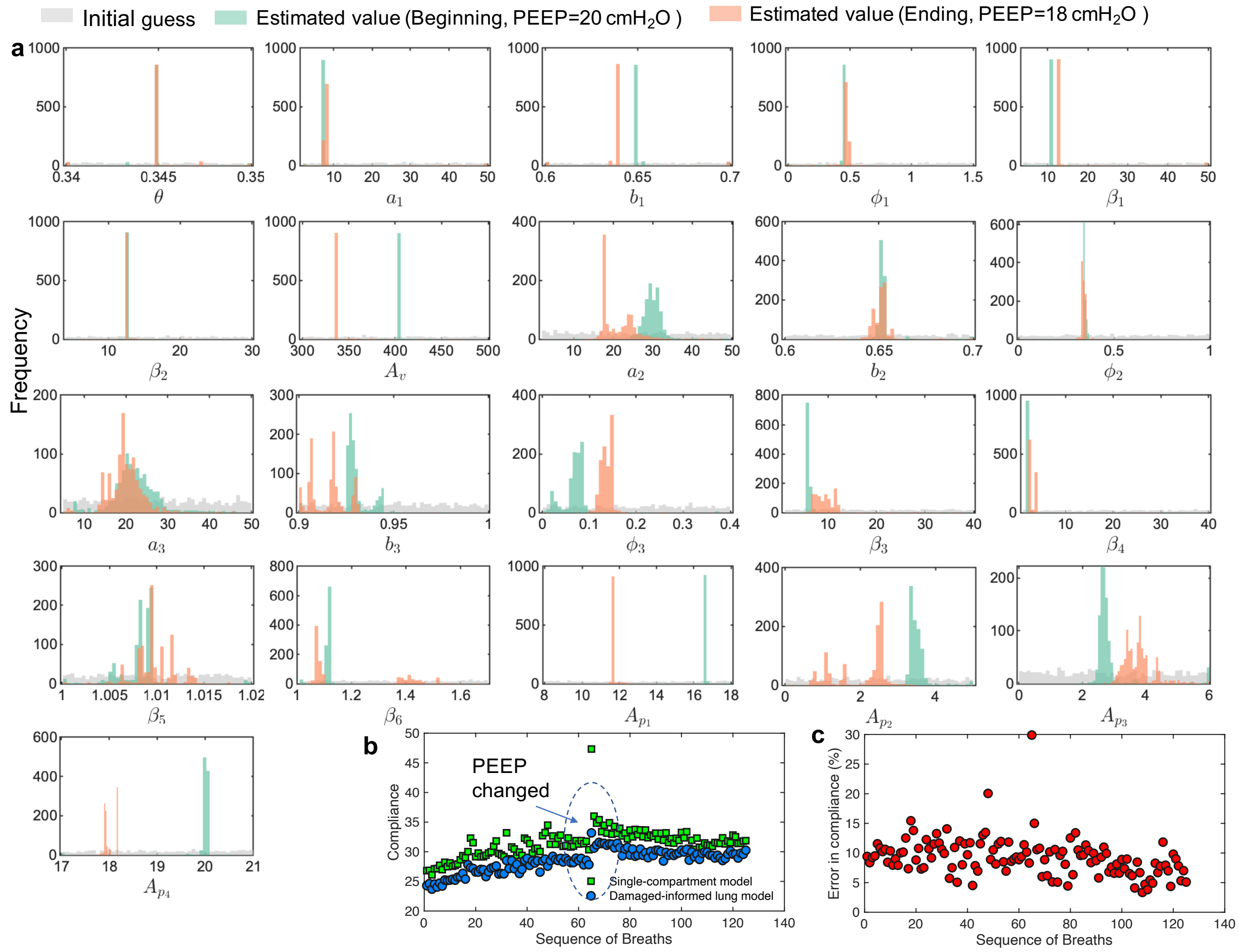


Figure S7: Histograms of the initial guesses and estimated parameters were obtained from 1000 samples that gave the lowest fitting error (see Methods). These results correspond to human patient 1 data shown in Fig. 6a. Note the variability in the reparatory frequency is due to human-triggered mode. (b) Comparing lung compliance values extracted by fitting the single-compartment model to the human data with the damaged-informed lung ventilator model. In the latter case, $A_v/A_{p_1}$ ratio was used to calculate the compliance using the parameters values shown in Table 1. (c) Percentage error in the single-compartment lung compliance values with respect to our model.

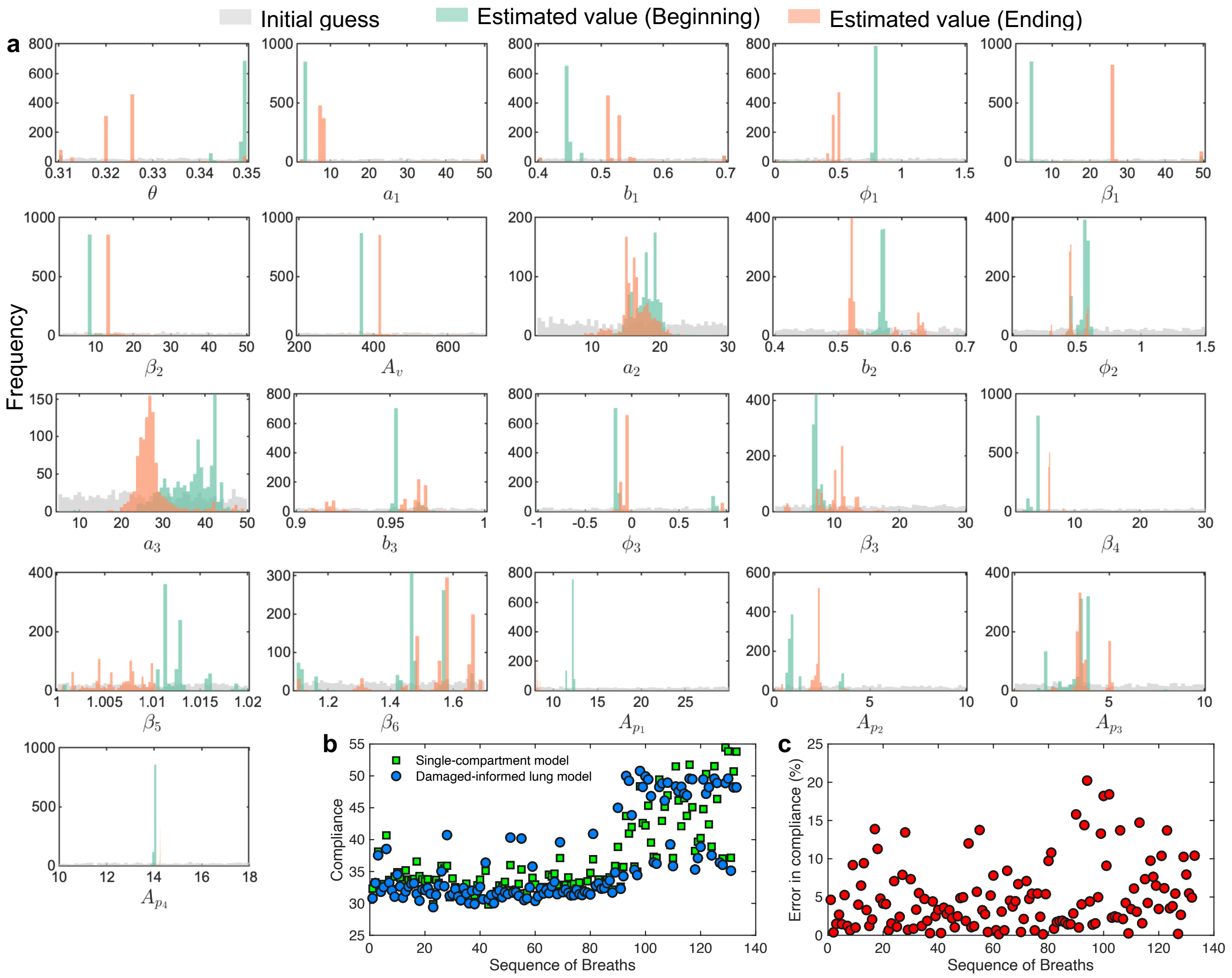


Figure S8: Histograms of the initial guesses and estimated parameters were obtained from 1000 samples that gave the lowest fitting error (see Methods). These results correspond to human patient 2 data shown in Fig. 6b. Note the variability in the reparatory frequency is due to human-triggered mode. (b) Comparing lung compliance values extracted by fitting the single-compartment model to the human data with the damaged-informed lung ventilator model. In the latter case, $A_v/A_{p_1}$ ratio was used to calculate the compliance using the parameters values shown in Table S1. (c) Percentage error in the single-compartment lung compliance values with respect to our model.

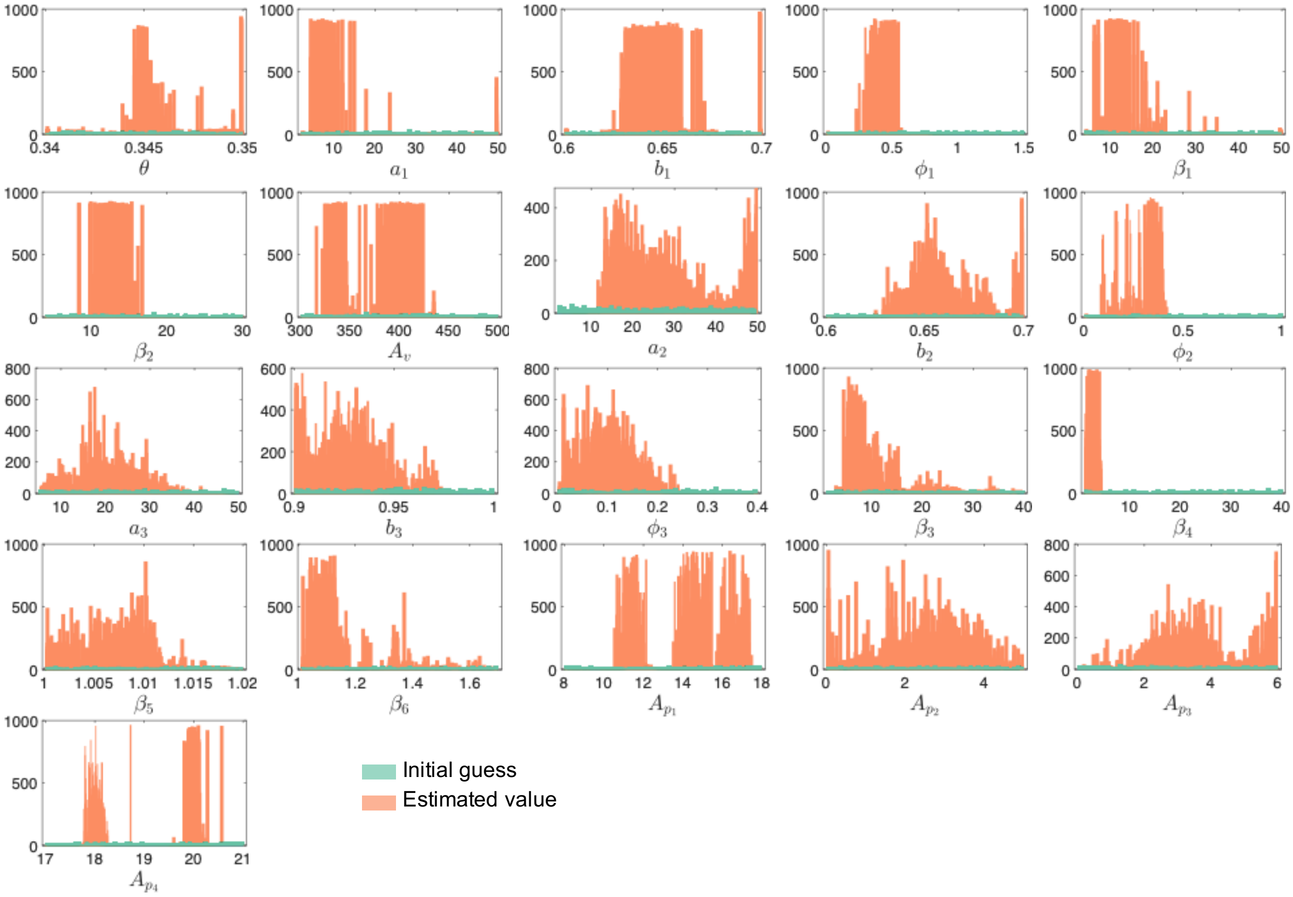


Figure S9: Histograms of the initial guesses and estimated parameters were obtained from 1000 samples that gave the lowest fitting error (see Methods). These results correspond to human patient 1 where of the thousands of breaths available, we selected a sequence of 130 breaths without VD at a random location. The PEEP value was changed from 18 to 20 $cmH_2O$ over the period. Note the variability in the reparatory frequency is due to human-triggered mode.

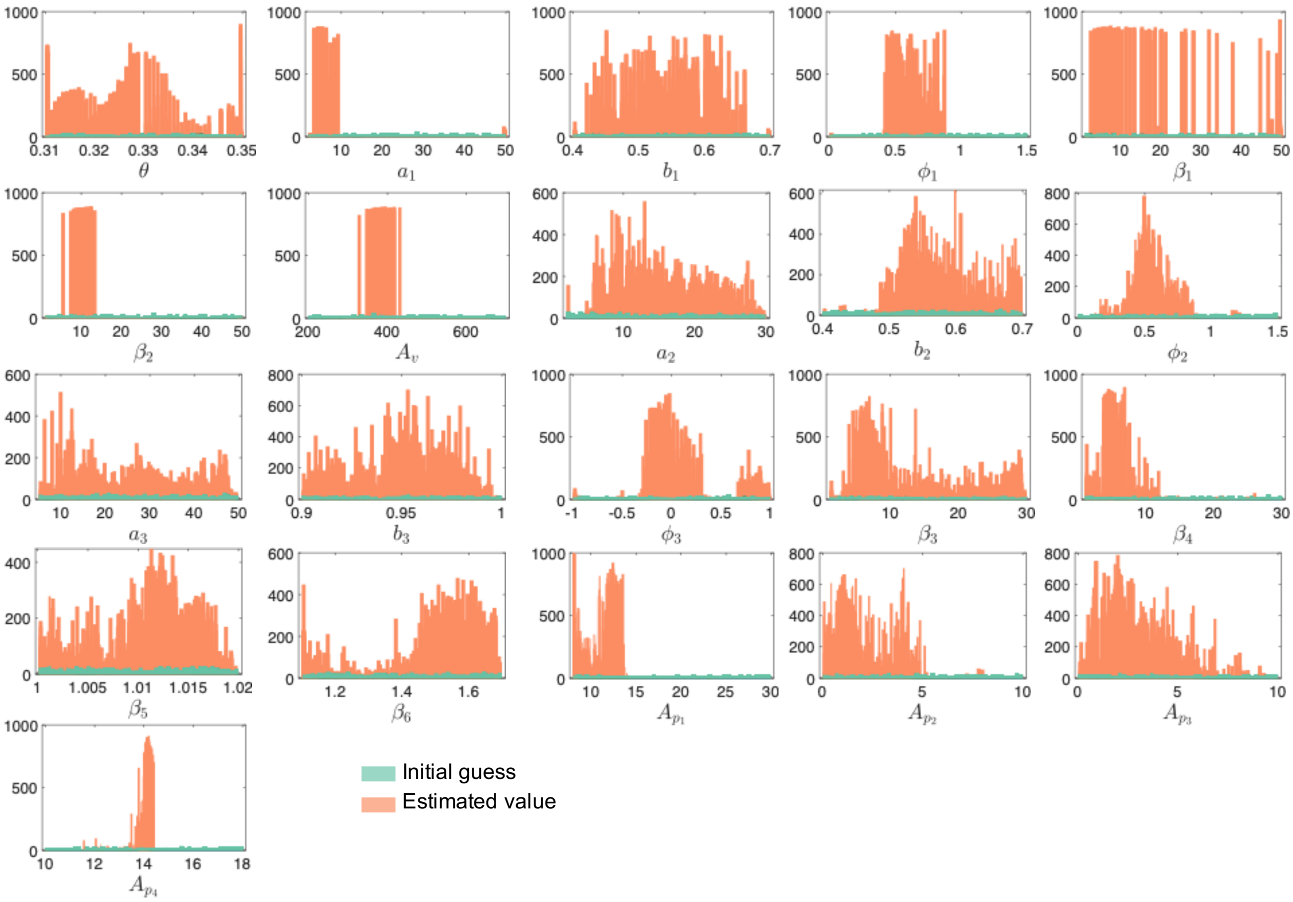


Figure S10: Histograms of the initial guesses and estimated parameters were obtained from 1000 samples that gave the lowest fitting error (see Methods). These results correspond human patient 2 where we randomly selected 133 breaths from the thousands of breaths observed. These breaths were selected from a several hours long time window where most of the breaths showed mild to severe VD. Note the variability in the reparatory frequency is due to human-triggered mode.
